# LRR protein RNH1 dampens the inflammasome activation and is associated with adverse clinical outcomes in COVID-19 patients

**DOI:** 10.1101/2021.04.12.438219

**Authors:** Giuseppe Bombaci, Mayuresh Anant Sarangdhar, Nicola Andina, Aubry Tardivel, Eric Chi-Wang Yu, Gillian M. Mackie, Matthew Pugh, Vedat Burak Ozan, Yara Banz, Thibaud Spinetti, Cedric Hirzel, Esther Youd, Joerg C. Schefold, Graham Taylor, Amiq Gazdhar, Nicolas Bonadies, Anne Angelillo-Scherrer, Pascal Schneider, Kendle M. Maslowski, Ramanjaneyulu Allam

## Abstract

Inflammasomes are cytosolic innate immune sensors of pathogen infection and cellular damage that induce caspase-1 mediated inflammation upon activation. Although inflammation is protective, uncontrolled excessive inflammation can cause inflammatory diseases and can be detrimental, such as in COVID-19. However, the underlying mechanisms that control inflammasome activation are incompletely understood. Here we report that the leucine rich repeat (LRR) protein Ribonuclease inhibitor (RNH1), which shares homology with LRRs of NLRP proteins, attenuates inflammasome activation. Deletion of RNH1 in macrophages increases IL-1β production and caspase-1 activation for inflammasome stimuli. Mechanistically, RNH1 decreases pro-IL-1β expression and induces proteasome-mediated caspase-1 degradation. Corroborating this, mouse models of monosodium urate (MSU)-induced peritonitis and LPS-induced endotoxemia, which are dependent on caspase-1, respectively show increased neutrophil infiltration and lethality in *Rnh1*^-/-^ mice compared to WT mice. Furthermore, RNH1 protein levels are negatively correlated with inflammation and disease severity in hospitalized COVID-19 patients. We propose that RNH1 is a new inflammasome regulator with relevance to COVID-19 severity.

## Introduction

The initiation of inflammation depends on pattern recognition receptors (PRRs), which sense invading pathogens or endogenous danger molecules that are released by dying cells. This process is critical in eliminating pathogens and initiating tissue repair (Gong et al., 2020). Inflammasomes are cytosolic PRRs that form multiprotein caspase-1-activating complexes upon activation to mediate inflammatory response. Inflammasome complexes contain sensors (e.g. NLRP3), an adaptor protein ASC (apoptosis-associated speck-like protein containing a caspase activation and recruitment domain) and effector protein caspase-1 (Martinon et al., 2009; Schroder and Tschopp, 2010). Assembly of these components into an inflammasome is initiated after sensing several host-derived danger-associated molecular patterns (DAMPs), bacterial toxins, nucleic acids, pathogenic crystals, and altered cellular components (Gong et al., 2020; Henao-Mejia et al., 2014). An active inflammasome complex catalyzes proteolytic cleavage of the pro-caspase-1 protein into functional caspase-1. Subsequently, caspase-1 processes IL-1 cytokine members pro-IL-1β and pro-IL-18 into biologically active IL-1β and IL-18 and initiates Gasdermin-D (GASMD) mediated pyroptosis, a form of cell death (Broz and Dixit, 2016). Several NOD-like receptor (NLRs) proteins (NLRP1, NLRP3, NLRP6, NLRC4/NAIP), HIN200 proteins (AIM2) and Pyrin can form inflammasome complexes (Schroder and Tschopp, 2010; Broz and Dixit, 2016). The NLR family is characterized by the presence of a central nucleotide-binding and oligomerization (NACHT) domain, which is commonly flanked by C-terminal leucine-rich repeats (LRRs) and N-terminal CARD or pyrin (PYD) domains. LRRs are believed to function in ligand sensing and autoregulation, whereas CARD and PYD domains mediate homotypic protein-protein interactions for downstream signaling (Schroder and Tschopp, 2010).

Ribonuclease inhibitor (RNH1) is a cytosolic LRR protein, however it can also be found in the nucleus and mitochondria (Dickson et al., 2005; Furia et al., 2011). The known function of RNH1 is to bind to pancreatic-type ribonucleases with femtomolar affinity and render them inactive (Dickson et al., 2005). In addition, RNH1 has been shown to inhibit oxidative damage, regulate angiogenin (ANG) mediated neovascularization and prevent tiRNA degradation (Monti et al., 2007; Dickson et al., 2009; Yamasaki et al., 2009). RNH1 has also been shown to bind and inhibit ER stress sensor IRE1 (Tavernier et al., 2018). However, the relevance of these observations *in vivo* has yet to be validated. We recently reported that RNH1 is important for embryonic erythropoiesis in mice, suggesting possible additional roles for RNH1 in mammalian physiology (Chennupati et al., 2018). RNH1 was the first cytosolic protein identified to contain LRRs (Dickson et al., 2005). LRRs are present in a large family of proteins that display vast surface areas to foster protein–protein and protein– ligand interactions (Kobe and Deisenhofer, 1995). LRR proteins are classified into subfamilies based on the organism of origin, cellular localization, LRR consensus sequence and length. To date, seven LRR subfamilies of proteins have been described: bacterial, ribonuclease inhibitor (RNH1)–like, cysteine-containing, SDS22, plant-specific, typical, and small (Dickson et al., 2005; Matsushima et al., 2010; Kobe and Kajava, 2001). The members of the RNH1-like subfamily are intracellular proteins and include human MHC class II transactivator (CIITA), Ran GTPase activating protein from *Saccharomyces pombe* (RANGAP1), and other NLRs (Dickson et al., 2005). Furthermore, the LRRs of RNH1 are very similar to those of NLRP proteins (Ng et al., 2011). However, whether RNH1 is involved in inflammasome activation has been not investigated.

Inflammasomes are critical to mount immune responses, however uncontrolled inflammasome activation is associated with several autoimmune and inflammatory diseases including Coronavirus disease 2019 (COVID-19) (Strowig et al., 2012; Berg and Velde, 2020). COVID-19 caused by severe acute respiratory syndrome coronavirus 2 (SARS-CoV-2) is associated with significant mortality and has resulted in more than 4 million deaths worldwide to date. The clinical manifestation of patients with COVID-19 includes pulmonary failure such as acute respiratory distress syndrome (ARDS) and cardiac injury (Huang et al., 2020a). Recent studies have reported increased inflammasome activation and inflammation in severe COVID-19 patients and suggest that increased inflammasome activation potentially mediates disease progression in COVID-19 (Lucas et al., 2020). Rodrigues et al., 2021; Kroemer et al., 2020; Toldo et al., 2020). It is therefore important to understand the molecules that control and resolve the overactivation of inflammasomes and inflammation in order to develop therapeutic targets for COVID-19 and other inflammatory disorders.

In this study, we found that RNH1 inhibits inflammasome activation by controlling proteasome-mediated degradation of the downstream effector molecule caspase-1. RNH1-deletion in mice displayed higher inflammasome-dependent activation of caspase-1. Interestingly, we also found that RNH1 expression in buffy coat and lung biopsies is negatively correlated with SARS-CoV-2-mediated inflammation and severity in COVID-19 patients. Collectively, these findings establish RNH1 as a previously unidentified negative regulator of inflammasome activation and indicates its potential role in human inflammatory diseases including SARS-CoV-2 mediated inflammation and pathology.

## Results

### RNH1 is predominantly expressed in myeloid cells and is increased under inflammatory conditions

Human RNH1 protein consists of 15 LRR repeats whose sequence and organization are similar to those present in multiple NLRP proteins (Ng et al., 2011; Hoss et al., 2019) (**Fig. 1A**). While NLR protein LRRs are confined to the C-terminal only, the RNH1 protein sequence consists entirely of LRRs and lacks any other identifiable functional domains (**Fig. 1B**). LRRs in NLRP family proteins are believed to function in ligand sensing and autoregulation (Schroder and Tschopp, 2010). However, whether RNH1 is also involved in inflammation similar to NLRP proteins has not been investigated. First, we checked RNH1 expression in haematopoietic cells by staining healthy human bone marrow biopsies for RNH1. We found that RNH1 is expressed at higher levels in myeloid cells compared to other hematopoietic cell-types such as lymphocytes and erythrocytes (**Fig. 1C**). Furthermore, RNH1 protein expression is elevated in bone marrow biopsies of patients with confirmed systemic inflammation (see methods) (**Fig. 1D**). Corroborating this, RNH1 protein levels were moderately increased in human monocyte cell-line (THP1 cells) upon stimulation with the TLR1/2 ligand Pam3Cysk4 (**Fig. 1E)** and in primary mouse bone marrow derived macrophages (BMDM) after the TLR4 stimulation with LPS **(Fig. 1F**). Collectively, these results suggest that RNH1 is predominantly expressed in myeloid cells and may play a role in inflammation.

**Fig 1.**
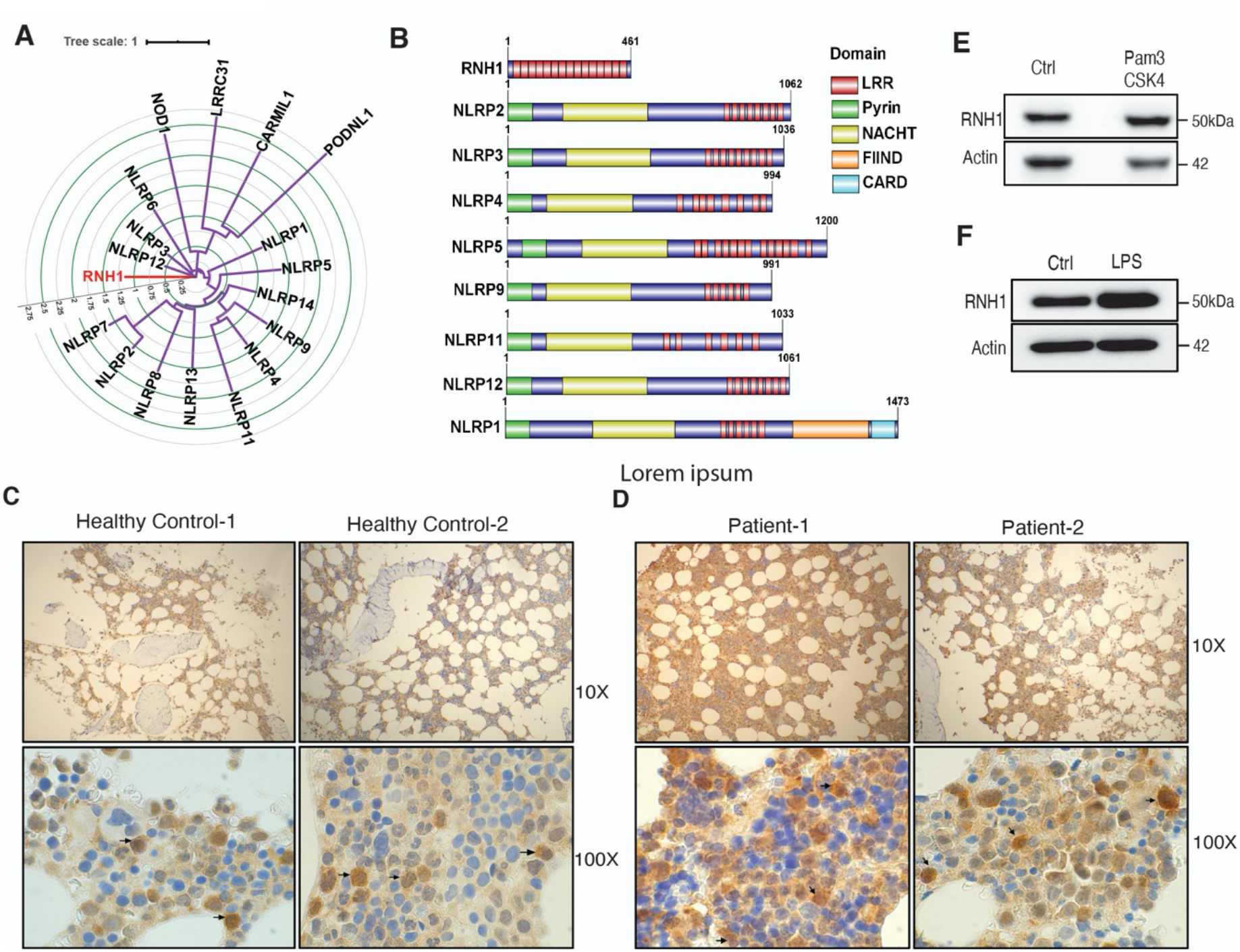
RNH1 shares homology with multiple NLR proteins and is expressed in myeloid cells. **(A)** Circular tree representing the domain conservation relationship of human RNH1. Protein sequence alignments were made using MAFFT. A maximum–likelihood phylogenetic tree was generated using IQ-Tree with 1000 bootstrap replicates. Internal tree scale is shown with circular grid. **(B)** Structural alignment of protein domains. The domain information of selected proteins taken from Uniprot and represented using Illustrator for Biological Sequences (IBS) tool. **(C)** Human healthy bone marrow biopsies were stained with RNH1 antibody. Myeloid cells showing high RNH1 expression are indicated with arrows. Original magnification 10X and 100X. **(D)** Bone marrow biopsies from patients with confirmed inflammatory conditions were stained with RNH1 antibody. Myeloid cells with increased RNH1 expression are indicated with arrows. Original magnification 10X and 100X. **(E)** THP1 cells were stimulated with TLR2 ligand Pam3CSK4 (1 µg/ml) for 24 hours. Total protein lysates were isolated and analyzed by Western blot with indicated antibodies. Blots are representative of two independent experiments. **(F)** Mouse primary BMDMs were stimulated with TLR4 ligand LPS (1 µg/ml) for 24 hours. Total protein lysates were isolated and analyzed by Western blot with indicated antibodies. Blots are representative of two independent experiments.

### Absence of RNH1 increases NLRP3 inflammasome activation

NLRP3 is the most well-studied inflammasomes and is activated by several pathogens and danger stimuli (Broz and Dixit, 2016). To investigate the role of RNH1 in NLRP3 inflammasome activation, we generated RNH1-deficient (RNH1-KO) THP1 cells using the CRISPR/cas9 system and stimulated them with NLRP3 agonists. RNH1-KO THP1 cells showed increased mature IL-1β production and caspase-1 cleavage compared to wildtype cells (WT) (**Fig. 2A and 2B**). Additionally, we also found increased pro-IL-1β and pro-caspase1 in unstimulated RNH1-KO cells compared to WT (**Fig. 2B)**. shRNA knockdown studies in THP1 cells also showed similar results (**Fig. S1A and B**). Since human and mouse RNH1 share 72.8% homology at the protein level, we also checked NLRP3 activation in mouse-derived immortalized macrophages (iMAC cells)(Broz et al., 2010). RNH1-KO iMAC cells also showed increased IL-1β production in response to NLRP3 agonists (**Fig. S2A**). This data suggests that RNH1 limits NLRP3 inflammasome activation in both human and mouse cells. Inflammasome activation also triggers pyroptosis, a type of inflammatory cell death (Miao et al., 2011). We indeed observed an increased trend of cell death in RNH1-KO cells compared to WT cells upon NLRP3 activation (**Fig. 2C and Fig. S2B**). In order to confirm the inhibitory function of RNH1 on inflammasome activation, we performed a transient reconstitution of RNH1 in RNH1-KO THP1 cells using lentiviral transduction, followed by stimulation with NLRP3 agonists. Reconstitution of RNH1 caused a decrease in IL-1β secretion (**Fig. 2D and E**). We also generated stable RNH1 over-expressing THP1 cells and stimulated them with NLRP3 agonists. RNH1 overexpression decreased IL-1β secretion, thereby confirming that RNH1 negatively regulates NLRP3 inflammasome activation (**Fig. 2F and 2G**). Furthermore, RNH1-KO THP1 cells also showed enhanced ASC speck formation upon nigericin stimulation, which is a direct indicator of inflammasome activation (Stutz et al., 2013) (**Fig. 2H and 2I**). Taken together, these results suggest that RNH1 dampens NLPR3 inflammasome activation.

**Fig 2.**
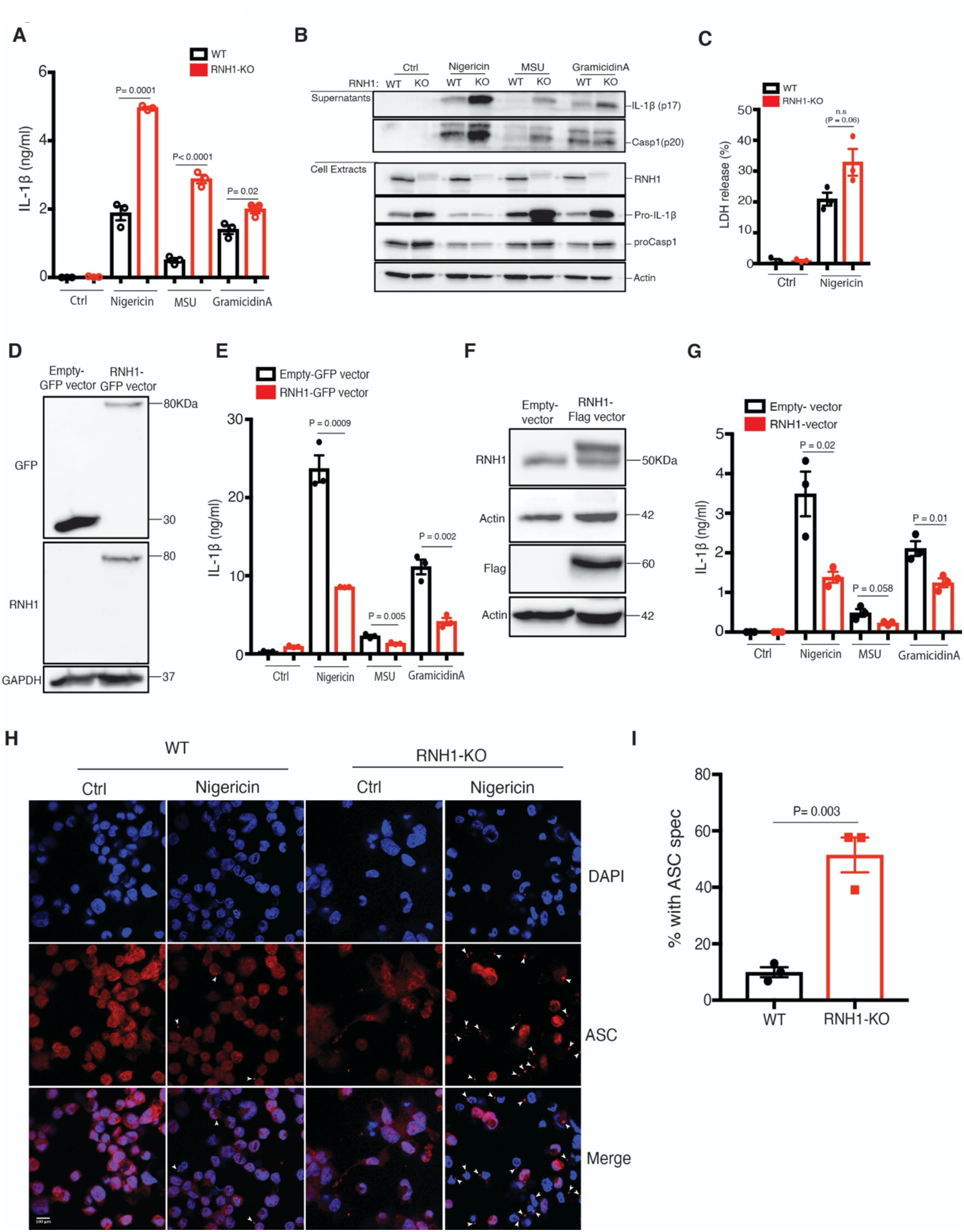
RNH1 inhibits NLRP3 inflammasome activation. **(A and B)** PMA differentiated wildtype (WT) and RNH1-KO THP1 cells were stimulated with Nigericin (5 µM) for 1h or with MSU (500 µg) and Gramicidin A (30 µg) for 5h. **(A)** Supernatants were collected and IL-1β ELISA was performed. Data are means ± SEM of pooled data from three independent experiments. Statistical analyses were performed using a two-tailed *t*-test. **(B**) Cell lysates and supernatants were analysed for pro- and cleaved-forms of caspase-1 and IL-1β by Western blot. Blots were representative of three independent experiments. **(C)** WT and RNH1-KO THP1 cells were stimulated with NLRP3 agonist Nigericin (5µM) for 18h. Supernatants were collected, and cell death was measured with LDH assay. Data are means ± SEM of pooled data from three independent experiments. Statistical analyses were performed using a two-tailed *t*-test. **(D and E)** RNH1-KO THP-1 cells transiently infected with GFP tagged control or RNH1 expressing lentivirus particles. **(D)** Cell lysates were analysed by Western blot to demonstrate the RNH1 reconstitution in RNH1-KO THP1 cells. Blots are representative of three independent experiments. **(E)** These cells were stimulated with Nigericin (5 µM) for 1h or with MSU (500 µg) or Gramicidin A (30 µg) for 5h. Supernatants were collected and IL-1β ELISA was performed. Data are means ± SEM of pooled data from three independent experiments. **(F and G)** Total cell lysates from THP1 cells constitutively expressing control or Flag-RNH1 were analysed by Western blot with indicated antibodies to demonstrate RNH1 overexpression **(F)**. Blots are representative of three independent experiments. These cells were stimulated with Nigericin (5 µM) for 1h or with MSU (500 µg) or Gramicidin A (30 µg) for 5h. Supernatants were collected and IL-1β ELISA was performed **(G)**. Data are means ± SEM of pooled data from three independent experiments. **(H and I)** Immunofluorescence microscopy analysis of ASC specks in THP1 cells stimulated with Nigericin (5 µM) for 1h. DNA staining is shown in blue (DAPI) and ASC staining is shown in red. Arrowheads indicate ASC inflammasome specks. (Scale bar: 100 μm) **(H)**. Quantification of ASC specks (**I**). Data are means ± SEM of pooled data analysing at least 10 fields from three independent experiments.

### RNH1 regulates NF-kB activation and is involved in the priming of NLRP3

Activation of the canonical NLRP3 inflammasome occurs in two steps (Latz et al., 2013). The priming step requires activation of nuclear factor kappa B (NF-κB), *e*.*g*. when LPS binds to TLR4, which subsequently upregulates NLRP3 and pro-IL-1 via activation of NF-κB. The second step requires a signal initiated by pathogen-associated molecular patterns (PAMPs) or DAMPs, which leads to NLRP3 activation and formation of an inflammasome complex with ASC and caspase-1. In RNH1-KO cells, we found increased levels of pro-IL-1β suggesting that RNH1 may also be involved in reducing the priming signal (**Fig. 2B**). To further understand this, WT and RNH1-KO THP1 cells were stimulated with the TLR2 ligand Pam3CSK4. RNH1-KO cells showed increased pro-IL-1β and NLRP3 expression in both a time- and Pam3CSK4 concentration-dependent manner, supporting the involvement of RNH1 in priming (**Fig. 3A and 3B**). The nuclear translocation and activity of NF-κB is tightly controlled by IκB - the inhibitor of NF-κB. Phosphorylation of IκB triggers its degradation and NF-κB activation. Interestingly, RNH1-KO cells also displayed decreased levels of IκB and increased levels of phospho-IκB compared to WT cells (**Fig. 3C**). To further confirm NF-κB activation, we monitored the production of pro-inflammatory cytokines and found that RNH1-KO cells exhibited enhanced secretion of TNF and IL-6 compared to WT cells (**Fig. 3D and 3E**). Collectively, these results suggest that RNH1 decreases NF-κB activation and therefore reduces the priming step.

**Fig 3.**
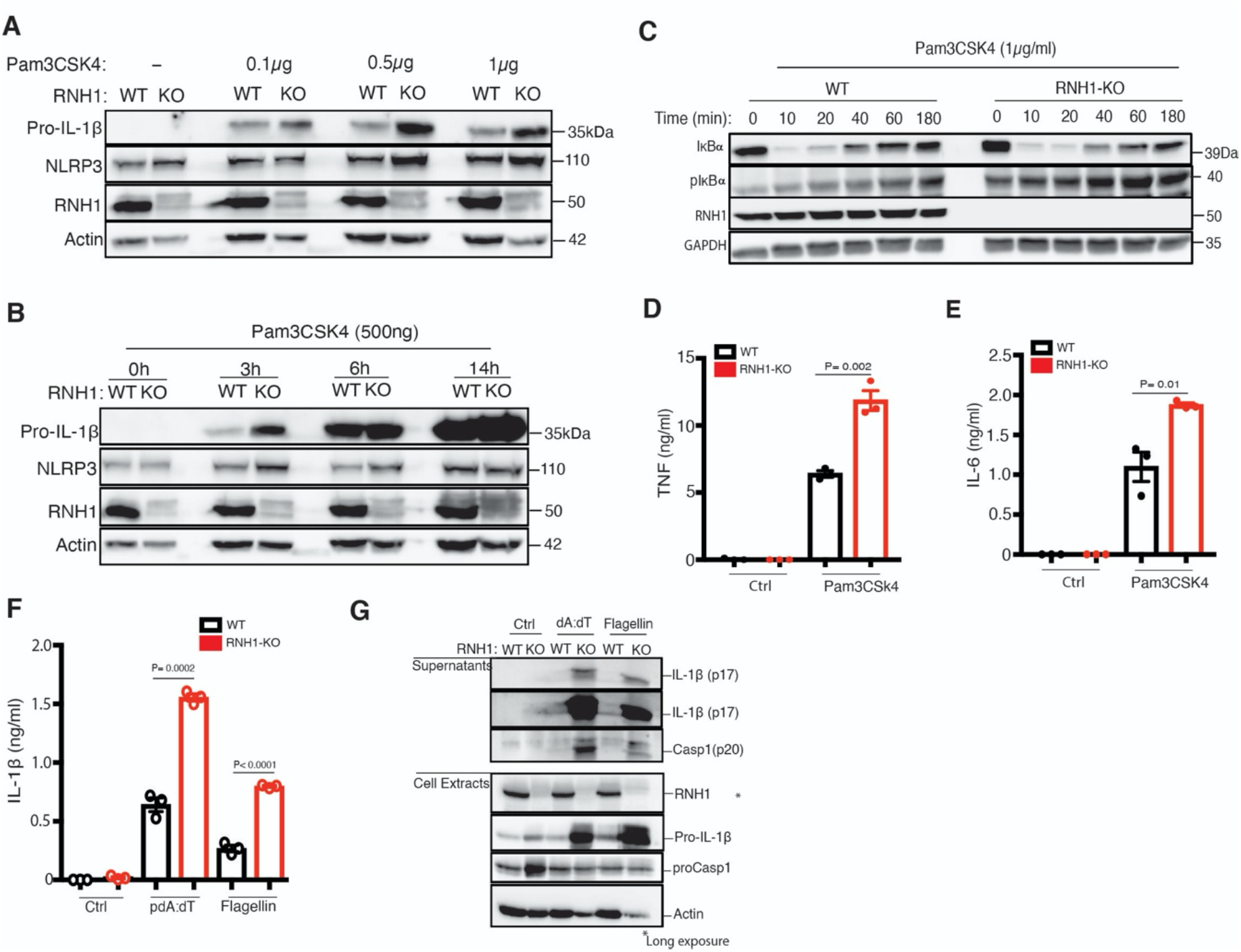
RNH1 negatively regulates activation of NF-κB, AIM2 and NAIP/NLRC4. **(A and B)** Undifferentiated WT and RNH1-KO THP1 cells were stimulated with TLR2 ligand Pam3CSK4 in a dose-(**A**) and time-(**B**) dependent manner as indicated. Total cell lysates analysed for pro-IL-1β and NLRP3 expression by Western blot. Blots are representative of three independent experiments. **(C)** Undifferentiated WT and RNH1-KO THP1 cells were stimulated with Pam3CSK4 (1µg/mL) for different time points as indicated. Total cell lysates analysed for NF-κB activation by Western blot with indicated antibodies. Blots are representative of three independent experiments. **(D and E)** Undifferentiated WT and RNH1-KO THP1 cells were stimulated with Pam3CSK4 (1µg/mL) for 6h and 18h. Supernatants were analyzed for TNF and IL-6 by ELISA, respectively. Data are means ± SEM of pooled data from three independent experiments. **(F and G)** PMA differentiated WT and RNH1-KO THP1 cells were transfected with AIM2 agonist poly dA:dT (5 µg) or NAIP/NLRC4 agonist cytosolic flagellin (600 ng) for 5h. Supernatants were collected and IL-1β ELISA was performed **(F)**. Data are means ± SEM of pooled data from three independent experiments. Statistical analyses were performed using a two-tailed *t*-test. Cell lysates and supernatants were analysed for pro- and cleaved-forms of caspase-1 and IL-1β by Western blot **(G)**. Blots are representative of three independent experiments.

### RNH1 deficiency increases AIM2 and NAIP/NLRC4 inflammasome activation

To check whether RNH1 is also involved in activation of AIM2 and NAIP/NLRC4 inflammasomes, we stimulated WT and RNH1-KO THP1 cells with their agonists. We used cytosolic dsDNA (pdA:dT) and flagellin to activate AIM2 and NAIP/NLRC4 inflammasomes, respectively. Interestingly, RNH1-KO showed an increase in mature IL-1β production and caspase-1 cleavage compared to WT cells (**Fig. 3F and G**), suggesting increased AIM2 and NAIP/NLRC4 inflammasome activation in RNH1-KO THP1 cells. We also found increased AIM2 inflammasome activation in mouse iMAC cells, however there was no significant increase in cytosolic flagellin-induced NAIP/NLRC4 inflammasome activation (**Fig. S2C**). Further studies are required to understand cell-type and species-specific differences in RNH1 mediated flagellin-induced inflammasome activation (Gram et al., 2021). To further confirm inflammasome activation, we checked ASC oligomerization by monitoring the abundance of ASC in the insoluble cell fraction (Lugrin and Martinon, 2017). As expected, the loss of RNH1 increased ASC oligomerization in response to NLRP3 and AIM2 inflammasome activators (**Fig. S3A**). We also found an increased trend of cell death in RNH1-KO cells by AIM2 activation (**Fig. S3B and C**). Together, these results suggest that RNH1 also negatively regulates AIM2 and NAIP/NLRC4 inflammasome activation.

### RNH1 mediates degradation of caspase-1 through the proteasome

The above results suggest that RNH1 inhibits not only NLRP3 inflmmasome but also AIM2 and NAIP/NLRC4 – possibly by acting on a common downstream effector/s. Interestingly, pro-caspase-1 protein levels were increased in RNH1-KO cells at steady state (**Fig. 4A and 2B**). Recently it has been reported that caspase-1 levels are regulated at the transcriptional level by NF-κB activation (Lee et al., 2015). However, qPCR analysis shows that expression of both *caspase-1* and *ASC* are equivalent in WT and RNH1 KO cells (**Fig. 4B**), indicating that RNH1 does not regulate caspase-1 at the transcriptional level. To check whether RNH1 regulates caspase-1 expression at the translational or post-translational level, we performed chase experiments using actinomycin-D, cycloheximide (CHX) and the proteasome inhibitor MG-132 in WT and RNH1*-*KO THP1 cells. Actinomycin-D and CHX inhibit transcription and translation respectively. Actinomycin-D and CHX treatment decreased caspase-1 protein levels in both WT and RNH1*-*KO cells with similar kinetics (**Fig. 4C and D**). Interestingly, proteasome blockade with MG-132 increased caspase-1 protein levels in WT cells. However, in RNH1*-*KO cells caspase-1 levels were not increased over time with MG-132 treatment. This suggests that loss of RNH1 might stabilize caspase-1 protein by inhibiting proteasome mediated degradation (**Fig. 4E**). To test this further, we expressed full length Flag-caspase-1 in the presence or absence of GFP-RNH1 in HEK293T cells. Strikingly, caspase-1 protein levels decreased with increasing concentration of RNH1, an effect that was blocked in the presence of MG-132 (**Fig. 4F**). It has been reported that overexpression of caspase-1 induces self-cleavage (Li et al., 2008), however, we did not observe caspase-1 self-cleavage at the concentration that we used in our experiments. Interestingly, we did not observe a similar decrease in another protease, OMI/HTRA2 (Walle et al., 2008) (**Fig. 4G**). This implies that RNH1 might specifically mediate proteasomal degradation of caspase-1 or of a defined set of proteins. Collectively, these results suggest that RNH1 increases proteasome-mediated caspase-1 degradation. However, further studies are required to understand the molecular mechanism of RNH1 mediated caspase-1 degradation.

**Fig 4.**
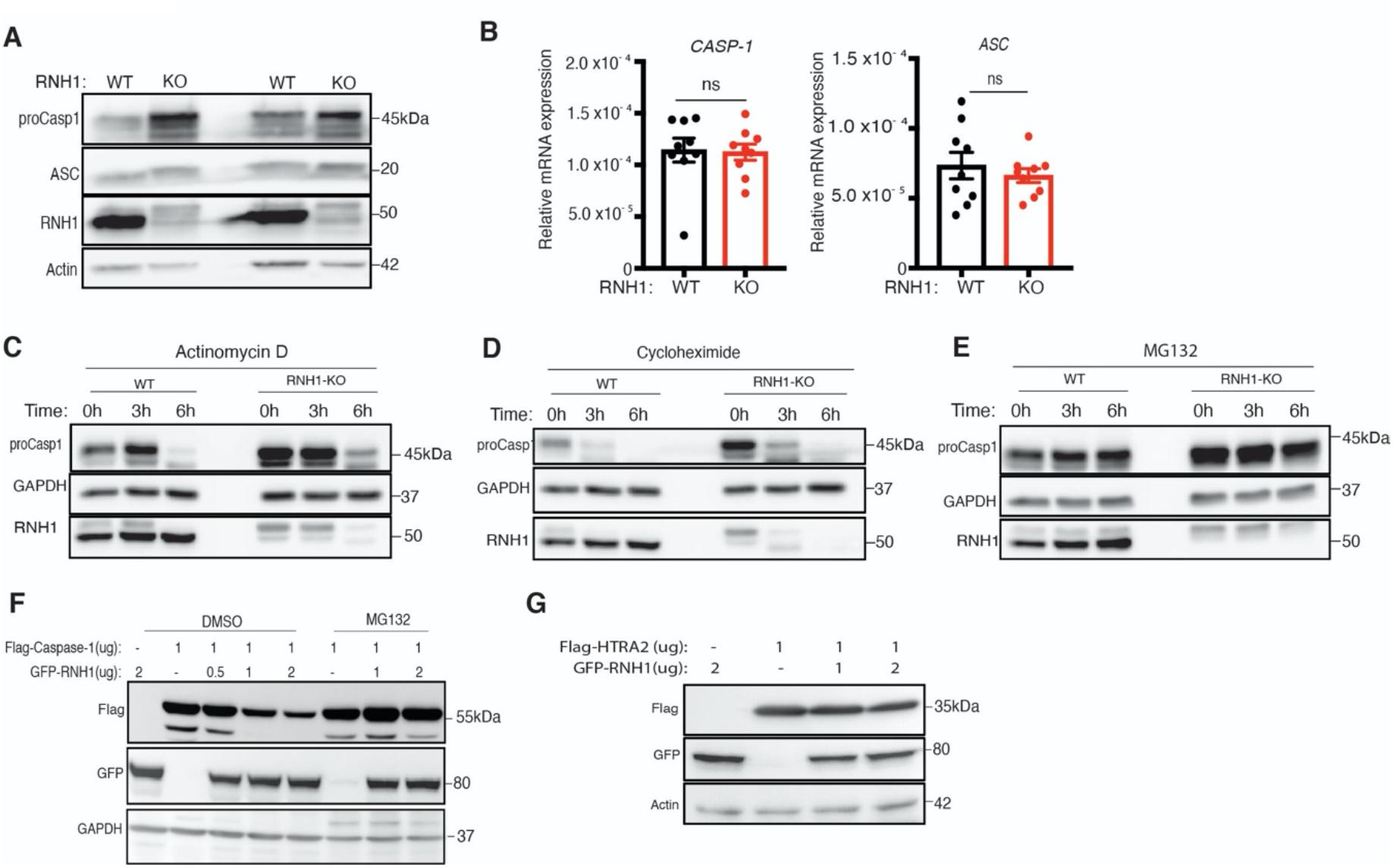
RNH1 increases caspase-1 degradation through the proteosome. **(A)** Total cell lysates from WT and RNH1-KO THP1 cells were analysed for pro-caspase-1 and ASC expression by Western blot. Blots are representative of two independent experiments. **(B)** qRT-PCR analysis for *CASP-1* and *ASC* mRNAs from WT and RNH1-KO THP1 cells. mRNA levels are normalized to 18S rRNA expression. Data shown as means ± SEM from three independent experiments. **(C-E)** WT and RNH1-KO THP1 cells were treated with Actinomycin D or Cycloheximide or with the proteasome inhibitor MG 132 for indicated time duration. Cell lysates were analysed for pro-caspase-1 protein levels by Western blot. Blots are representative of three independent experiments. **(F)** HEK293T cells were treated with or without MG-132 and transfected with Flag-tagged casapase-1 with different concentration of GFP-tagged RNH1 plasmid as indicated. Cells were harvested and cell lysates were analyzed for caspase-1 by Western blot with the indicated antibodies. Blots are representative of three independent experiments. **(G)** HEK293T cells were transfected with Flag-tagged OMI/HTRA2 plasmid with different concentrations of GFP-tagged RNH1 plasmid as indicated. Cells were harvested and cell lysates were analyzed by Western blot with indicated antibodies. Blots are representative of three independent experiments.

### Loss of RNH1 in mice increases inflammation and lethality upon inflammasome activation

Constitutive deletion of *Rnh1* leads to embryonic lethality in mice (Chennupati et al., 2018). We generated *Rnh1* conditional knockout mice (*Rnh1*^*fl/fl*^) to understand the *in vivo* relevance of our results (**Fig. 5A**). *Rnh1*^*fl/fl*^ mice were crossed with transgenic interferon-inducible *Mx1-Cre* mice to generate an inducible *Rnh1*^*-/-*^ mouse model (*Rnh1* ^*fl/f;Mx1-Tg+*^). Administration of polyinosinic:polycytidylic acid (polyI:C) into *Rnh1* ^*fl/f;Mx1-Tg+*^ mice leads to cre-recombination and complete deletion of RNH1 protein expression in hematopoietic organs such as the bone marrow (BM), which we refer to as *Rnh1*^*-/-*^ mice (**Fig. 5B and C**). We generated BM-derived macrophages (BMDM) from WT (*Rnh1*^*fl/fl*^) and *Rnh1*^*-/-*^ mice and then stimulated them with inflammasome activators. As observed in cell lines, loss of RNH1 in primary mouse BMDMs led to increased mature IL-1β production and caspase-1 cleavage compared to WT cells upon stimulation with NLRP3, AIM2 and NAIP/NLRC4 agonists (**Fig. 5D and E**). To investigate the *in vivo* role of RNH1 in inflammasome activation, we performed mouse models of caspase-1-dependent MSU-induced peritonitis and LPS-induced lethality (Martinon et al., 2006; Li et al., 1995; Man et al., 2017) (**Fig. 5F**). Since *Mx1-Cre* activation also deletes genes in non-haematopoietic organs (Kühn et al., 1995), we transplanted WT (*Rnh1*^*fl/fl*^) and *Rnh1*^*-/-*^ (*Rnh1*^*fl/fl*^ *Mx1-Cre*^***+***^) BM into irradiated CD45.1 congenic mice to exclude *Rnh1* deletion effects from non-hematopoietic cells. After 8 weeks, we checked reconstitution levels in WT and *Rnh1*^*-/-*^ BM transplanted CD45.1 congenic mice. We found comparable reconstitution in WT and *Rnh1*^*-/-*^ BM transplanted CD45.1 congenic mice via FACS analysis (**Fig. 5G**). *Rnh1* was deleted by giving three rounds of 200 μg of polyI:C injections. Four days after the last injection, mice were analyzed for peripheral blood parameters. At this time point, we observed no major difference in leukocyte numbers between WT and *Rnh1*^*-/-*^ BM transplanted CD45.1 congenic mice (**Fig. 5H**). In a MSU-induced model of peritonitis, *Rnh1*^*-/-*^ mice showed increased neutrophil infiltration in the peritoneal cavity compared to WT mice (**Fig. 5I**). Consistent with this, we also found increased IL-1β production in the peritoneal lavage of *Rnh1*^*-/-*^ mice compared to WT mice (**Fig. 5J**). We next challenged mice with LPS at 10 mg/kg. This dose was lethal to *Rnh1*^*-/-*^ mice but all WT mice survived (**Fig. 5K**). Altogether, these results demonstrate that *Rnh1* deficiency promotes excessive inflammasome activation and increases inflammation *in vivo*.

**Fig 5.**
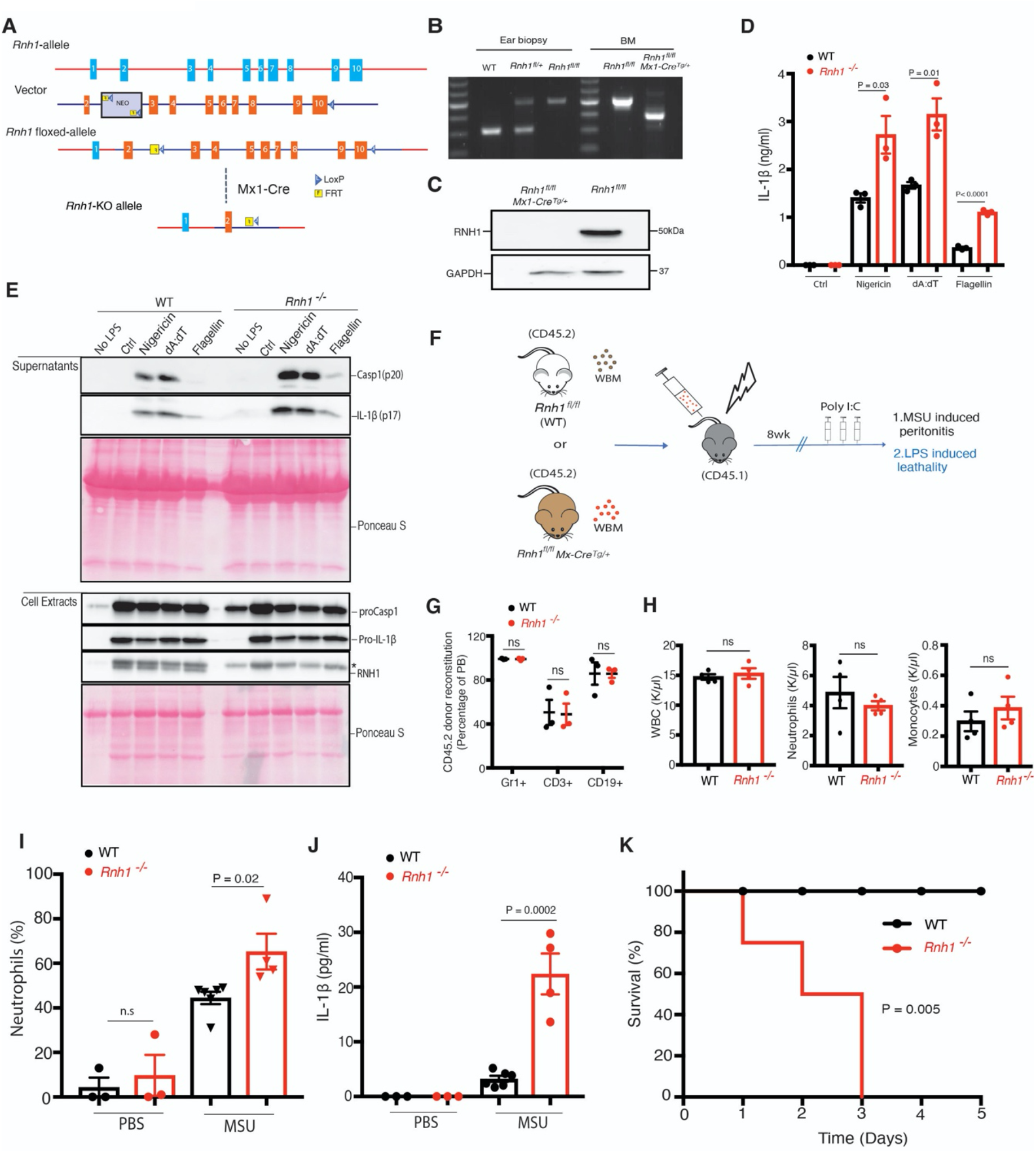
*Rnh1* deficiency promotes inflammation in mice. **(A)** Schematic showing design of *Rnh1-*floxed targeting vector (see methods for details) **(B)** DNA isolated from mouse ear biopsies was genotyped by PCR. Primers were designed to distinguish WT and floxed allele sequence. 314 bp size corresponds to WT, and 514 bp size corresponds to floxed gene. To detect *Rnh1* deletion after Cre-recombination in *Rnh1*^*fl/fl*^ MxCre^+^ mice (after one week of polyIC treatment), a third primer was used which amplifies 365 bp size (see methods). **(C)** Total protein lysates from BM cells of WT and *Rnh1*^*-/*^ mice were analyzed by Western blot with the indicated antibodies. Blots are representative of 3 independent experiments. **(D and E)** BMDMs from WT and *Rnh1*^*-/-*^ mice primed with LPS (100 ng) for 3h and then cells were stimulated with Nigericin (5 µM) for 1h or transfected with poly dA:dT (5 µg) or flagellin (600 ng) for 5h. **(D)** Supernatants were collected and IL-1β ELISA was performed. Data are means ± SEM and representative of three independent experiments. Statistical analyses were performed using a two-tailed *t*-test. **(E)** Cell lysates and supernatants were analysed for pro- and cleaved-forms of caspase-1 and IL-1β by Western blot. Blots were representative of two independent experiments. **(F)** Schematic showing *in vivo* experimental setup. Bone marrow was transplanted from WT (*Rnh1*^*fl/fl*^) or *Rnh1*^*-/-*^ *(Rnh1*^*fl/fl*^ MxCre^+^) (CD45.2) into irradiated recipients (CD45.1). After 8 weeks of reconstitution mice were treated with Poly(I:C) (300 μg/mouse) once every two days for three doses. Four days later mice were used for the MSU peritonitis and LPS endotoxemia lethality models. **(G)** Post transplantation reconstitution levels were monitored after 8 weeks in the peripheral blood (PB) as indicated (n = 3 mice). Data are shown as mean ± SEM. *P* values were determined by 2way ANOVA with Sidak’s multiple comparisons test. **(H)** PB counts of WBC, neutrophils and monocytes in WT and *Rnh1*^*-/-*^ mice one week after polyIC injections. Data are shown are mean ± SEM. *P* values were determined by a two-tailed *t*-test (n = 3). **(I and J)** WT and *Rnh1*^*-/-*^ mice received intraperitoneal injection of MSU (1 mg/mouse) or sterile PBS. After 12h, peritoneal lavage fluid was taken and neutrophils infiltration analysed by FACS **(I)**. Peritoneal lavage supernatants were collected and IL-1β levels were quantified by ELISA **(J)**. Data shown as means ± SEM and *P* values were determined by a two-tailed *t*-test (n = 3-6 per group). **(K)** Kaplan–Meier survival curves of WT and *Rnh1*^*-/-*^ mice after LPS (10 mg/kg) treatment (n = 4 mice).

### RNH1 is negatively associated with disease severity in COVID-19 patients

Recent studies suggest that pronounced inflammation and inflammasome activation in patients with COVID-19 correlates with increased disease severity and poor prognosis (Berg and Velde, 2020; Mehta et al., 2020; Jose and Manuel, 2020; Toldo et al., 2020; Rodrigues et al., 2021; Kroemer et al., 2020; Ferreira et al., 2021). Since our results suggest that RNH1 negatively regulates NF B and inflammasome activation, we investigated RNH1 levels in COVID-19 patients. Buffy coat samples from critically ill COVID-19 patients admitted to the intensive care unit (ICU) (n= 17) and COVID-19 patients admitted to general COVID-19 ward (n=11 patients) were analyzed for RNH1 protein expression by Western blot (see methods for details). Disease severity of these patients is established by comorbidity data and routine clinical parameters such as C-reactive protein (**Table. S1**). Strikingly, COVID-19 patients in ICU had significantly less RNH1 expression when compared to COVID-19 patients in general ward (**Fig. 6A, B and Fig. S4**). The decreased RNH1 expression in patients admitted to ICU is unlikely to reflect leukocyte numbers, since leukocyte numbers were rather increased in ICU patients compared to general ward COVID-19 patients (**Table. S1**). To further support these findings, we analyzed RNH1 expression in lung sections of patients from an independent study in the UK who deceased either from COVID-19 (n = 8) or from non-viral causes (n =13) (**Table. S2**). In line with the data above, RNH1 expression is largely absent in the lungs of deceased COVID-19 patients, whereas patients who succumbed to non-viral causes did show RNH1 staining in infiltrating cells in the lung (**Fig. 6C and Fig. S5**). This data illustrates a loss of RNH1 expression in immune cells during severe COVID-19, which may suggest a crucial role for RNH1 in controlling severe inflammation.

**Fig 6.**
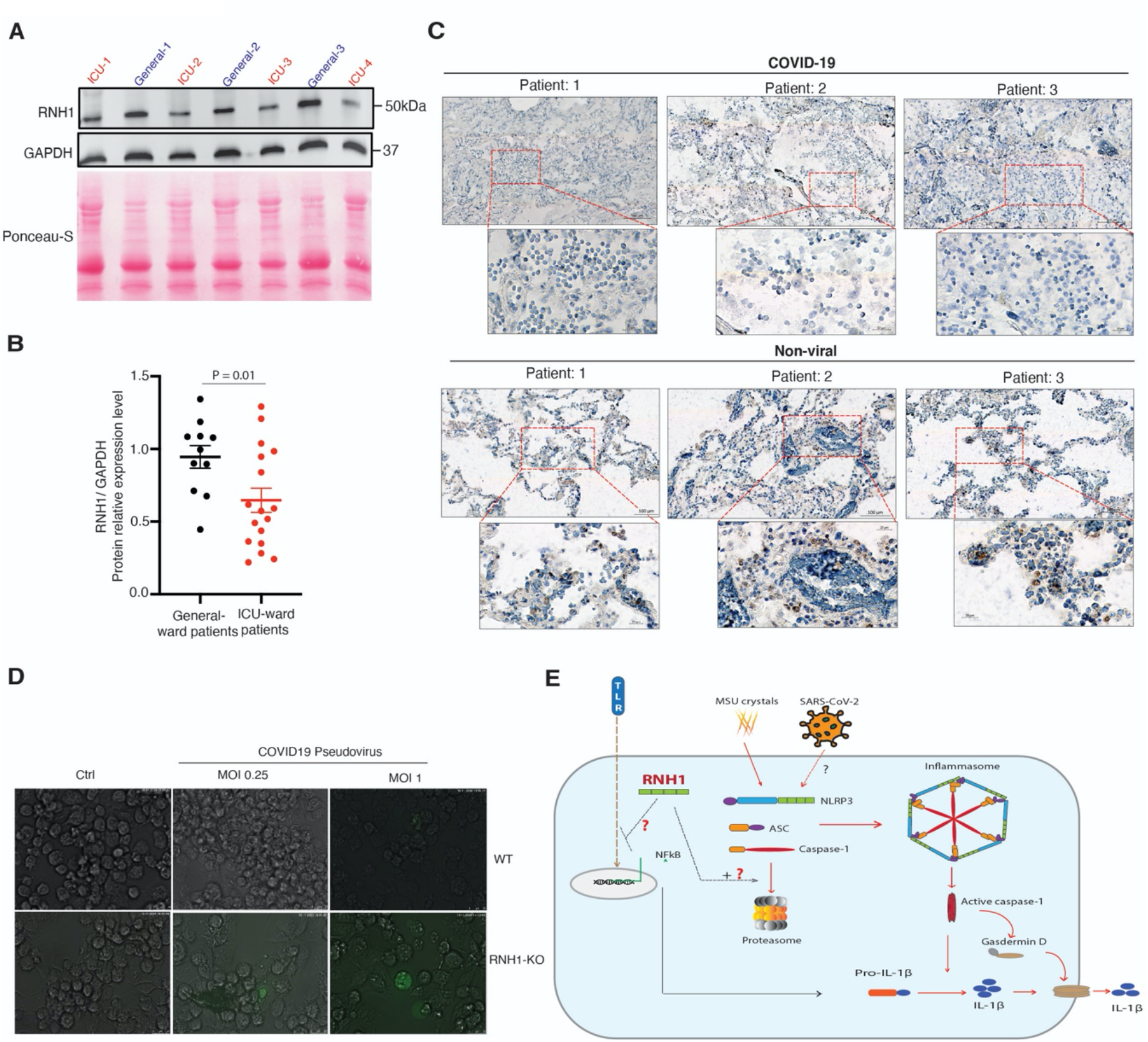
Decreased RNH1 expression correlates with disease severity in COVID-19. **(A)** Total cell lysates from peripheral blood buffy coats of ICU and general ward admitted COVID-19 patients were analysed for RNH1 protein levels by Western blot. Blots were repeated three times. **(B)** RNH1 protein levels from blots were quantified by ImageJ analysis for each patient and normalized with respective GAPDH protein levels. **(C)** Postmortem lung tissue from deceased persons with either COVID-19 or non-viral causes of death were stained for RNH1 and imaged using a Zeiss axioscan Z1. Subsequent image analysis was performed using Zeiss ZEN software to extract images at different magnifications. Insets show higher magnification of area indicated in the red boxes. Brown staining indicates RNH1 positive cells. **(D)** PMA differentiated WT and RNH1-KO THP1 cells were infected with SARS-CoV-2 pseudovirus for 24 hours. Infection efficiency was monitored by measuring GFP signal with fluorescence microscopy. Images are representative of two independent experiments. (Scale bar 25 µm). **(E)** Illustration of RNH1 mediated anti-inflammatory mechanisms. First, RNH1 could potentially inhibit NF-κB signaling through an unknown mechanism. Second, RNH1 regulates inflammasome activation by controlling caspase-1 protein levels via proteasome-mediated degradation.

Recent studies document the critical role of monocytes and macrophages in SARS-CoV-2 infection (Ferreira et al., 2021; Desai et al., 2020). To check whether RNH1 affects the SARS-CoV-2 infection rate in macrophages, we infected WT and RNH1-KO macrophages differentiated from THP1 monocytes with SARS-CoV-2 pseudoviral particles (Hoffmann et al., 2020; Nie et al., 2020). These pseudoviral particles express SARS-CoV-2 spike protein together with enhanced GFP (eGFP) as a reporter. Interestingly, we found increased eGFP+ cells in RNH1-KO cells compared to WT cells (**Fig. 6D**). At a multiplicity of infection (MOI) of 0.25, we observed 1% of RNH1-KO cells were positive for eGFP cells. This increased to 2.5% at an MOI of 1, however we did not find any eGFP+ cells in the WT at 0.25 MOI and very few (<0.5%) at 1 MOI. These results suggest that RNH1-KO cells are more susceptible to infection (**Fig. 6D**). Collectively, our findings suggest that decreased expression of RNH1 strongly correlates with severity and increased inflammation in COVID-19 patients, and possibly increases the risk of SARS-CoV-2 infection.

## Discussion

Inflammasomes are multiprotein cytoplasmic complexes that induce potent inflammation and Gasdermin-D-mediated pyroptosis upon activation (Broz and Dixit, 2016). This process is critical for pathogen clearance and to maintain tissue homeostasis. However, uncontrolled inflammasome activation leads to excessive inflammation and tissue destruction, which is a primary cause of several inflammatory, autoimmune and autoinflammatory diseases (Strowig et al., 2012; Lamkanfi and Dixit, 2011). Therefore, precise regulation of inflammasome signaling is necessary to prevent collateral damage while still preventing pathogen insurgence. Our results unravel a new mechanism involving RNH1 acting as a negative regulator of inflammasome activation. Conventionally, RNH1 is known to inhibit ribonucleases and protect RNA from degradation (Dickson et al., 2005). However, constitutive deletion of *Rnh1* in mice is embryonically lethal because of marked defects in erythropoiesis. These defects originate from decreased translation of the transcriptional erythropoiesis regulator GATA-1 rather than detectable effects on mRNA levels. This suggests that RNH1 fulfills functions beyond inhibition of ribonucleases (Chennupati et al., 2018). This is also supported by the present study in which RNH1 mainly exerts post-translational effects.

The RNH1 protein sequence consists of only LRRs and evolved via exon duplication (Haigis et al., 2002). Interestingly, the LRRs of RNH1 share homology with the LRRs of NLRs, yet the function of RNH1 in NLR signaling is unknown. Although RNH1 is a ubiquitously expressed protein (Dickson et al., 2005), we demonstrate that it is highly expressed in myeloid cells and its expression is increased under inflammatory conditions, thus pointing to a potential role in regulating inflammation. Interestingly, we found that RNH1-deleted macrophages have increased IL-1β production, caspase-1 activation and pyroptosis in response to NLRP3 agonists, suggesting that RNH1 can potentially inhibit NLRP3-induced inflammation. Increased expression of RNH1 in inflammatory conditions may be a compensatory mechanism to dampen inflammation. Additionally, we identified increased NF-κB signaling and increased expression of pro-IL-1 and NLRP3 in the absence of RNH1. This clearly indicates that RNH1 regulates NLRP3 inflammasome activation at the priming step as well as during the activating signal. We did not investigate mitogen-activated protein kinase (MAPK) pathways, which play a role in TLR induced inflammation (Arthur and Ley, 2013), and therefore cannot exclude that the MAPK pathway may mediate priming in RNH1-deleted cells. Previous reports suggest that reactive oxygen species (ROS) are involved both in priming and in NLRP3 inflammasome activation (Zhou et al., 2010). RNH1 protein contains numerous cysteine residues (e.g. 32 in human RNH1), whose sulfhydryl groups may play a structural role and have also been shown to protect against oxidative damage (Dickson et al., 2005). Thus, increased ROS production in the absence of RNH1 may be responsible for NLRP3 inflammasome activation. However, loss of RNH1 also increased AIM2 and NAIP/NLRC4 inflammasome activation, both of which do not require ROS. Therefore, these observations do not substantiate a major anti-ROS role of RNH1 to suppress activation of the NLRP3 inflammasome. Further studies are necessary to exclude the role of ROS in RNH1-mediated NLRP3 inflammasome regulation. From this study, RNH1 has emerged as an inhibitor for inflammasome activation. By suppressing inflammasome activation, RNH1 likely controls the extent of potentially dangerous immune activation.

In addition to increased canonical inflammasome activation and higher caspase-1 activity in RNH1-KO cells, we also observed increased caspase-1 protein levels in RNH1-KO cells compared to WT. Caspase-1 is a cysteine protease acting downstream of inflammasome activation. Upon activation, pro-caspase-1 is cleaved into two subunits (p10 and p20) and mediates proteolytic processing of pro-IL-1β and pro-IL-18 while promoting pyroptosis by cleaving gasdermin (Broz and Dixit, 2016). Few studies report mechanisms for the inhibition of caspase-1 function. Flightless-I binding can inhibit caspase-1 cleavage (Li et al., 2008; Man et al., 2017) and CARD only protein -1 (COP-1) has been shown to interact with the CARD domain of caspase-1 and inhibit its interaction with ASC (Druilhe et al., 2001; Pedraza-Alva et al., 2015). However, to our knowledge, regulation of caspase-1 expression at post-translational level as found in the present study has not been previously reported. Furthermore, we demonstrate that RNH1 promotes proteasome-mediated caspase-1 protein degradation, which was rescued by inhibiting proteasome function. Further investigation is required to understand how RNH1 regulates proteasome-mediated caspase-1 degradation. Caspase-1-deficient mice are resistant to LPS-induced lethality and MSU-induced peritonitis (Martinon et al., 2006; Man et al., 2017). Supporting a role for RNH1 in inflammasome activation and caspase-1 activity, *Rnh1*^*-/-*^ mice showed markedly increased susceptibility to LPS-induced lethality. Furthermore, *Rnh1*^*-/-*^ mice also displayed increased neutrophil infiltration in MSU-induced peritonitis. These results mirror observations obtained *in vitro*. Collectively, our study unravels a significant role of RNH1 in dampening inflammasome activation and demonstrates a previously unidentified regulation of caspase-1 through proteasome-mediated degradation (**Fig. 6E**).

Uncontrolled inflammation mediates immune pathology in several inflammatory diseases including pandemic COVID-19 (Mehta et al., 2020; Jose and Manuel, 2020; Huang et al., 2020b). Recent studies have demonstrated a correlation of increased inflammasome activation and disease severity in COVID-19 patients (Berg and Velde, 2020; Toldo et al., 2020). We have showed that RNH1-KO THP1 cells are susceptible to infection with SARS-CoV-2 pseudo virus. Furthermore, we identified a significant decrease in RNH1 protein expression in severe ICU-admitted COVID-19 patients compared to less severe general ward-admitted COVID-19 patients, suggesting that RNH1 expression negatively correlates with COVID-19 disease severity. Additionally, these results were confirmed in an independent COVID-19 patient cohort using post-mortem lung tissue. We showed that RNH1 protein expression is decreased in lung sections of COVID-19 patients compared to patients who died from non-viral causes. This is contrary to what we have shown in the systemic inflammatory conditions, where increased RNH1 expression may be a compensatory mechanism to inhibit inflammation (**Fig. 1D**). Therefore, it is unclear what drives the reduced expression of RNH1 in severe COVID-19 pathology. Understanding whether this is a common feature of severe inflammation, or of infection-driven inflammation, and what causes it is of interest.

The data provided here using animal models of inflammation, as well as patients with COVID-19 pathology strongly support a role for RNH1 in suppressing lethal inflmammation warrenting further investigation into the mechanisms. Increasing our understadning could pave the way for development of RNH1 as a prognostic tool and/or development of small molecules to enhance RNH1 expression or function.

## Acknowledgements

We thank Fabio Martinon, University of Lausanne for helpful discussions. We thank Alexandra Moniz for critical reading of the manuscript. We thank the FACS and microscopy facility of the DBMR (University of Bern) for assistance in flow cytometry and confocal experiments, respectively. This work was supported by the Swiss National Science Foundation (PP00P3_157486, PP00P3_183721 & PP00P3_190073) to RA.

## Author contributions

GB, MAS, NA, KMM, CHY performed experimental work. AT maintained mouse lines and helped with experimental work. NB and YB helped with histopathology of human bone marrow samples.AG and VBO helped with COVID-19 pseudovirus infection studies. TS, CH and JCS provided COVID-19 patients samples and data. KMM, GM, MP, EY and GT helped with histopathology of COVID-19 lung biopsy data from UK. AAS and PS helped with reagents and advice. RA designed, supervised the research work and wrote the manuscript with inputs from all the authors.

## Materials and methods

### Details of Rnh1 conditional-knockout mice generation and mouse experiments

*Rnh1* conditional-knockout (*Rnh1*^*fl/fl*^) mice were generated on a C57BL/6 background. A 13.8 kb region used to construct the targeting vector was first subcloned from a positively identified C57BL/6 BAC clone (RP23: 217N5) using a homologous recombination-based technique. The region was designed in a way that the long homology arm (LA) extends 6.23 kb 3’ to the single LoxP site. The short homology arm (SA) extends about 3.0 kb 5’ to the LoxP/FRT flanked Neo cassette. The single LoxP site is inserted downstream of exon 10, and the Neo cassette is inserted upstream of exon 3. The size of the target region is about 4.6 kb containing exons 3-10 (**Fig. 5A**). *Rnh1* targeting vector was electroporated into (C57BL/6) embryonic stem (ES) cells. Targeted (C57BL/6 FLP) embryonic stem cells were microinjected into Balb/c blastocysts. Resulting chimeras with a high percentage black coat colour were mated to C57BL/6 WT mice to generate Neo deleted, F1 heterozygous offspring. Germline transmission was confirmed by PCR of tail genomic DNA. Screening of *Rnh1*^*fl/fl*^ mice by PCR genotyping was carried out using the following primers on ear genomic DNA: 5’-ACATGGTGTTCTGGG TGTACGGTGG-3’ (forward in the intron region after second exon), 5’-CTGAGTAAGGAC TGCTGGGCTGAG-3’ (reverse in the proximal *LoxP* region). This reaction amplifies 314 bp in size for WT allele and 514 bp size for floxed allele (**Fig. 5B**). We crossed *Rnh1*^*fl/fl*^ mice with *Mx1*-*Cre* mouse strain (Jackson: 002527) to generate inducible mouse model (*Rnh1*^*fl/fl*^ *Mx1-Cre*^***+***^). To exclude non-haematopoietic deletion, total bone marrow (BM) was isolated from WT (*Rnh1*^*fl/fl*^) or *Rnh1*^*fl/fl*^ *Mx1-Cre*^***+***^ mice and transplanted (7×10^6^ BM cells) into lethally irradiated CD45.1^+^ congenic recipient mice (8 weeks old male mice) (B6.SJL-*Ptprc*^*a*^*Pepc*^*b*^/BoyCrl, from Charles river). After 8 weeks of reconstitution, *Rnh1* was excised by giving three rounds of 200 μg poly(I:C) (Invivogen) using intraperitoneal injections once in two days. We used these mice for MSU induced peritonitis model and LPS lethality model. To detect *Rnh1* deletion after Cre-recombination, we used a third primer 5’-AAGACCCATCCAGAGCCGAGG-3’ (reverse in the distal intron region) in above mentioned PCR reaction using DNA form BM cells, which amplifies 365 bp size (**Fig. 5B**). Western blot was also performed in BM cells to check RNH1 protein expression (**Fig. 5C**). Mouse peripheral blood was taken from the lateral tail vein with an EDTA coated Microvette 100 K3E (Sarstedt, 20.1278). Full blood counts were measured on an IDEXX ProCyte Dx™ Hematology Analyzer (IDEXX Laboratories). The genotypes of the mice could not be blinded or randomized, due to the experimental design. All groups of mice (experimental and control mice) were age- and sex-matched. We used littermates for our experiments. All animal experiments were approved by the Swiss Federal Veterinary Office (Bern, Switzerland) under valid authorization (BE39/16). Mice were handled according to Swiss Federal Veterinary Office guidelines under valid authorization. Mouse strains are available upon request.

### MSU-induced peritonitis model

Mice were injected intraperitoneally (IP) with sterile PBS or MSU (1 mg/mouse) to induce peritonitis. After 12 h mice were euthanized by CO2-asphyxiation and the peritoneal cavity was flushed with sterile PBS. The lavage fluid was centrifuged and supernatants were analyzed by ELISA for IL-1β production and the pelleted cells were analyzed for the influx of neutrophils (CD45^+^, Ly-6G^+^, CD11b^+^) by BD LSRII flow cytometer (BD Biosciences). Data was analysed by using FlowJo (version 9.3.1, TreeStar Inc) software.

### LPS-induced septic shock

Mice were injected intraperitoneally (IP) with LPS (10 mg/kg) or PBS and survival rates were monitored.

### Histopathology and bone marrow biopsies

Healthy human controls and inflammatory disease patients bone marrow biopsies were obtained from Swiss MDS Registry. Biopsies were stained with human RNH1 antibody from Prestige Antibodies^®^ Sigma (HPA039223). These studies were approved by the competent local human ethics committee (ID:2017-00734).

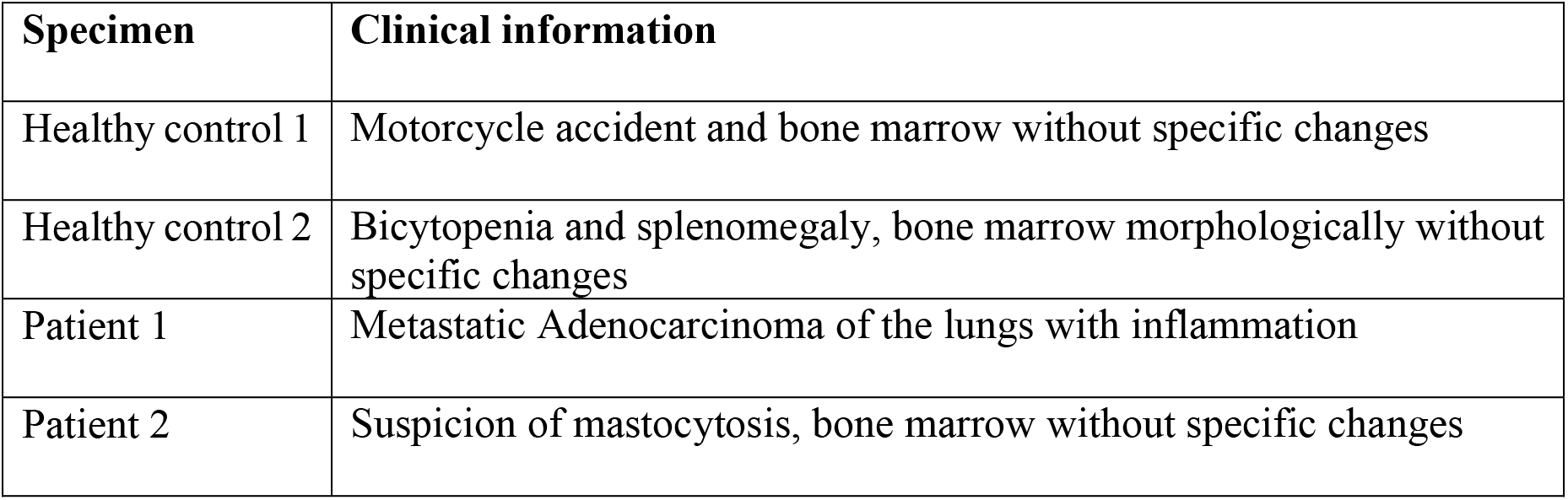

### Cells and cell culture media

Mouse bone marrow derived macrophages (BMDM), iMACs (immortalized macrophages) and Human THP-1 cells were used for inflammasome activation experiments.THP-1 cells and iMACs were grown in RPMI 1640 GlutaMAX™-I medium (Invitrogen) were supplemented with 10% (vol/vol) FBS (Amimed) and 1% of penicillin and streptomycin (PAA Laboratories), and incubated at 37 °C and 5% CO2. BMDMs were generated by established protocols. Briefly, total bone marrow cells were isolated from the femur and tibias of mice and differentiated into macrophages for 7 days in complete DMEM medium (500 mL, ThermoFisher) supplemented with M-CSF (40 ng/mL) (315-02, Peprotech).

### Reagents and plasmids

We purchased Nigericin (N7143), pdA:dT (poly(dA-dT)•poly(dT-dA)) sodium salt (P0883), Gramicidin A (G5002), PMA (P1585), Actinomycin D and Cycloheximide from Sigma-Aldrich, Ultra-pure flagellin (AG-40B-0025) from Adipogen, ultrapure LPS (LPS-EK), MSU (tlrl-msu) and Pam3Cysk4 (tlrl-pms) from Invivogen.

Cloning of full-length caspase-1 and OMI were amplified by PCR and sub-cloned into the mammalian expression vectors pCR3 in frame with the N-terminal flag tags. Full-length RNH1-Flag construct was described previously (Chennupati et al., 2018). To construct full length GFP-RNH1 plasmid, a human RNH1-full coding sequence (ORIGEN RC 200082) insert was cloned into the entry vector pENTR4-GFP-C1 (W392-1, Addgene) and then recombined into the destination vector pLenti CMV Blast DEST (706-1, Addgene) plasmid, using the Gateway cloning method.

### Stimulation experiments

All cells were stimulated at a density of 1×10^6^ cells per ml as describe previously (Allam et al., 2014). For inflammasome studies, THP1 cells were treated with PMA (100ng/ml) to differentiate into macrophages or BMDMs and iMACs were pre-stimulated for 3 hours with 100 ng/mL of ultrapure LPS. Later, cells were stimulated with 1 hour with 5uM of Nigericin or 5µg/mL of pdA:dT, 30µg/mL of Gramicidin A, 300µg/mL MSU and 200ng/mL of Flagellin. pdA:dT and flagellin were transfected with Lipofectamine 2000 according to the manufacturer’s protocol (Invitrogen). For all conditions, cell-free supernatants were analyzed by ELISA for cytokine secretion or cells were lysed for immunoblot analysis.

### CRISPR/CAS9-mediated RNH1-knockout THP1 and iMAC cell line generation

RNH1-KO THP1 cells (human monocytic cell line) were generated as described previously (Chennupati et al., 2018). For RNH1-KO iMAC cells CRISPR sequences targeting exon 2 (RNH1-KO-1) and exon 3 (RNH1-KO-2) of mouse RNH1 were designed and obtained KO cells as described previously (Chennupati et al., 2018). Target exon and the seed sequences preceding the protospacer adjacent motif (PAM) are the following: RNH1-1 oligo 1-5′-*CACC*GTCTGA TCCAGCAATACGAAG-*3’*; RNH1-1oligo 2—5′-AAACCTTCGTATTGCTGGA TCAGAC-3′; RNH1-2 oligo 1-5′-*CACC*GGATAACCCTATGGGGGACG*C*-*3* ′; RNH1-2 oligo 2-5′-AAACGCGTCCCCCATAGGGTTATCC-3’. All generated THP1 and iMAC clones were tested negative for mycoplasma contamination using MycoAlert Mycoplasma Detection Kit (Lonza, Cat#LT07-318). THP1 cells were obtained from ATCC, and iMAC cells were provided by Petr Broz (University of Lausanne) (Broz et al., 2010).

### Generation of stable THP1 cells expressing Flag–RNH1

Flag-RNH1 was further sub-cloned into retroviral vector pMSCVpuro (Clontech). Retroviral vector pMSCVpuro-Flag–RNH1 was co-transfected with the helper plasmids VSV-G and Hit60 into HEK293T cells using PEI transfection reagent. Culture supernatants containing recombinant viral particles were harvested and used to infect THP1 cells. To establish stable cell lines, THP1 cells were selected with puromycin (5 µg ml^-1^) three days after infection.

### Generation of transient THP1 cells expressing GFP–RNH1

RNH1-KO THP1 cells were infected with lentiviruses expressing RNH1 (pLenti CMV Blast GFP-RNH1) or the empty vector pLentiCMV-GFP-Blast (659-1, Addgene) as previously described (Papin et al., 2007). After 48 h post-infection, efficiency was monitored by GFP expression, and blasticidin (5 μg/mL) selected THP1 cells were plated for inflammasome assay.

### Generation of stable THP1 cells expressing shRNA-RNH1

Various lentiviral shRNA plasmids against RNH1 were purchased from Sigma and lentvirus was generated as previously described (Chennupati et al., 2018). To establish stable cell lines, THP1 cells were selected with puromycin (5 µg ml^-1^) three days after infection.

### Immunoblot analysis

Precipitated media supernatants or cell extracts were analyzed by standard immunoblot techniques. The following antibodies were used: anti-human RNH1 (A9), anti-mouse RNH1 (C10), anti-human ASC (Sc-514414), anti-GAPDH (G9, sc-365062) from Santa Cruz Biotechnology; anti- human IL-1β antibody (12242), anti-NFκB2 (p100), anti-IκBα, anti-Phospho-IκBα, anti-GFP (2555) from Cell signaling; anti-human caspase-1 (p20, Bally-1), anti-mouse caspase-1 (p20, AG-20B-0042), anti-NLRP3 (Cryo-2), anti-mouse ASC (AG-25B-0006) from Adipogen; anti-mouse IL-1β antibody (AF-401-NA) from R&D; anti-β-Actin from Abcam.

### TLR priming and NF-κB activation

WT and RNH1-KOTHP1 cells were stimulated with TLR2 agonist Pam3CSK4 (500ng/mL) at different time points 3, 6 and 14h or at different doses 100, 500 and 1000 ng/mL for 3h. For NF-*κB* activation, cells were stimulated at 10, 20, 40, 60 and 180 minutes by Pam3CSK4 (1000 ng/mL). After stimulation cell lysates were isolated and analysed via Western blot.

### Caspase-1 expression in HEK293T cells

HEK293T cells were treated with or without MG-132 and transfected with full length flag-tagged casapase-1 or OMI/HTAR2 plasmids with or without GFP-tagged RNH1 plasmid with the indicated concentration. At 24 hour, cells were harvested and cell lysates were analyzed by immunoblot. HEK293T cells routenly tested negative for mycoplasma contamination using MycoAlert Mycoplasma Detection Kit (Lonza, Cat#LT07-318). HEK293T cells were obtained from ATCC.

### Cytokine measurement and LDH assay

Cell supernatants were analyzed for cytokines and Lactate Dehydrogenase (LDH). Human and mouse IL-1β, TNF and IL-6 cytokine secretion measured by ELISA according to the manufacturer’s instructions (eBioscience). LDH was measured by colorimetric NAD linked assay by in-house developed kit according OPS Diagnostics instructions.

### Detection of ASC oligomerization by Western blot

We detected ASC oligomerization by Western blot as described previously (Lugrin and Martinon, 2017). Briefly, WT and RNH1-KO THP1 cells were primed with 100 ng of PMA and treated with Nigericin (5 μM) for 1 hour and Poly(dA:dT) (5 μg), for 5 hours. After stimulation, cells were detached and lysed on ice cold lysis buffer (20 mM Hepes·KOH, pH 7.5, 10 mM KCl, 1.5 mM MgCl2, 1 mM EDTA, 1 mM EGTA and 320 mM sucrose) by syringing 35 times using a 21G needle. Cell lysates were centrifuged for 8 min at 1,800 *x g* and supernatants were collected (30 μL of the supernatants were kept aside to check ASC expression in the lysates). Supernatants were further diluted two times with lysis buffer and then centrifuged at 2,000 *x g* for 5 min. Next, supernatants were diluted with 1 vol of CHAPS buffer (20 mM Hepes·KOH, pH 7.5, 5 mM MgCl2, 0.5 mM EGTA, 0.1 mM PMSF and 0.1% CHAPS) and then centrifuged for 8 min at 5,000 *x g*. The supernatant was discarded and the pellet was resuspended in 50 μL of CHAPS buffer supplemented with 4 mM of disuccinimidyl suberate (Thermo Scientific) and incubated at room temperature for 30 min. After incubation, samples were centrifuged for 8 min at 5,000 *x g*, and pellets were resuspended in SDS sample buffer without reducing agents and Western blot was performed.

### Detection of ASC spec formation by confocal microscopy

To further confirm the ASC specks formation, we performed confocal microscopy as described previously (Micco et al., 2016). Briefly, PMA-primed WT and RNH1-KO THP1 cells were seeded on coverslips in 12-well plates before inflammasome activation. Next day, cells were stimulated for 1h with Nigericin (5 μM) and fixed with 4% (vol/vol) PFA (Applichem) for 20 min at room temperature (RT). After fixation, cells were washed 3 times with 1x PBS for 5 min and permeabilized with 0.1% Triton X-100 in 1x PBS for 15 min at RT. Cell were washed with 1x PBS and blocked with 4%(w/vol) BSA for 30 min at RT. After blocking, cells were incubated overnight at 4 °C with the primary antibodies against ASC (HASC-71, BioLegend). Cells were washed 3 times with washing buffer (1x PBS and 0.025% of Tween20) for 5 min and incubated for 1h at RT with the Alexa Fluor secondary antibodies (Thermo Scientific). Cells were washed twice with washing buffer for 5 min and incubated in with DAPI (1 μg/mL) for 1 min. Again, cells were washed and coverslips were mounted on glass slides using mounting media (ProLong® Gold Antifade). Images were captured with inverted confocal microscope LSM 710 (Zeiss) and analyzed with ImageJ across at least 10 fields from three independent experiments.

### Actinomycin D, Cycloheximide and MG-132 chase assays

To study caspase-1 mRNA and protein stability, WT and RNH1-KO THP1 cells were treated with the following inhibitors for 0, 3 and 6 hours. Cell were treated with Actinomycin D (10 μg/mL), Cycloheximide (75 μg/mL), proteasomal inhibitor MG 132 (20 μM) to assess transcriptional, translational and posttranslational changes. After treatment, cells were harvested and cell lysates were analyzed by immunoblot.

### RNA preparation and qRT-PCR

Total RNA was isolated from WT and RNH1-KO THP1 cells using the QIAGEN RNeasy Kit according to the manufacturer’s protocol. Complementary DNA was generated from total RNA as described previously (Allam et al., 2015). The SYBR Green Dye detection system was used for quantitative real-time PCR on Light Cycler 480 (Roche). Controls consisting of ddH2O were negative for target and housekeeping genes. The following gene-specific primers (Microsynth) were used. 18SrRNA:5’-GCAATTATTCCCCATGAACG-3’(f), 5’-AGGGCCTCACTAAACCATCC-3’ (r); ASC: 5’-CCT CAG TCG GCA GCC AAG-3’ (f), 5’-CAG CAG CCA CTC AAC GTT TG-3’ (r); CASP1: 5’-TCC TCA GGC TCA GAA GGG AA-3’ (f), 5’-TGC GGC TTG ACT TGT CCA T-3’ (r); RNH1: 5’-GATCTGGGAGTGTGGCATCA-3’ (f), 5’-CTGCAGGACTTCACCCACAG-3’ (r).

### COVID-19 pseudovirus infection

The SARS-CoV-2 pseudoviral particles expressing COVID-19 spike protein pGBW m4137384: S protein was purchased from Addgene (149543) and the virus particles were produced as describe previously (Hoffmann et al., 2020; Nie et al., 2020). WT and RNH1-KO THP1 cells were treated with PMA (100 ng/mL) to differentiate into macrophages and infected with SARS-CoV-2 pseudovirus at MOI of 20 and 100 for 24 hours. Infection efficiency was monitored by measuring GFP signal using fluorescent microscope (Leica DMI 400 B).

### COVID-19 patient samples

We analysed a total of 28 COVID-19 patients samples from a previously published study (n= 13) (Spinetti et al., 2020), but with addition of new patients to the cohort (n=15). We isolated protein samples from buffy coats of COVID-19 patients and analysed RNH1 expression by Western blot. The present study is part of a larger COVID-project (NCT04510012) and data of some patients were previously investigated regarding immune functionality in COVID-19 (Spinetti et al., 2020). The study was performed in accordance with the “Declaration of Helsinki” and approved by the Kantonale Ethikkommission KEK, Bern, Switzerland, Nr. 2020-00877.

### COVID-19 tissue microarray samples

Post-mortem (PM) tissues were collected as part of the ethically approved LoST-SoCC study in accordance with the “Declaration of Helsinki” and Human Tissue Authority legislation. The study was approved by the Newcastle North Tyneside 1 research ethics committee 19/NE/0336, IRAS 193937. Non-COVID causes of death were all non-respiratory, non-infectious, obtained from PMs that were done prior to the pandemic, and lungs were morphologically within normal limits. The original FFPE tissue blocks were taken from representative samples at post-mortem, formalin fixed and processed as per standard diagnostic blocks. For tissue microarray (TMA) construction, representative 3mm tissue cores were cored out of FFPE tissue blocks and re-blocked within a 4×4 format TMA, sectioned and mounted onto glass slides.Slides were dehydrated and endogenous peroxides quenched. Antigen retrieval was done using TRIS buffer pH9 boiled for 15 minutes. Slides were incubated with permeabilization buffer (0.3% Triton-X100 in PBS) then blocking buffer (10% FBS in PBS). Anti-RNH1 (Prestige Antibodies^®^ Sigma, HPA039223) was added overnight at 1:500, followed by anti-rabbit secondary, ABC amplification and DAB detection (both ThermoFisher). Slides were then counterstained with Meyer’s haematoxylin and coverslips applied with aqueous mounting agent and slides imaged using Zeiss axioscan Z1. Images were analysed using Zen Lite software (Zeiss).

### Sequence conservation analysis

Human RNH1 protein sequence was used as a query to perform Blastp with human refseq proteins. From Blast output, selected protein sequences (excluding the isoforms and model refseq proteins) were aligned using MAFFT multiple sequence aligner (Katoh and Standley, 2013). Sequence conservation analysis was performed on IQ-Tree webserver (http://iqtree.cibiv.univie.ac.at/) using ultrafast bootstrap analysis with 1,000 number of bootstrap alignment (Katoh and Standley, 2013). Circular tree was visualized using Interactive Tree Of Life (iTOL) (Letunic and Bork, 2019). For structural alignment of protein domains, the domain information of selected proteins was taken from Uniprot and represented using Illustrator for Biological Sequences (IBS) tool (Liu et al., 2015)Sequences used for multiple sequence alignment: NP_002930.2_RNH1; NP_653288.1_NLRP12; NP_789792.1_NLRP14; NP_703148.4_NLRP5; NP_001120933.2_NLRP3; NP_659444.2_NLRP11; NP_789790.2_NLRP9; NP_001167553.1_NLRP2; NP_604393.2_NLRP4; NP_789780.2_NLRP13; NP_001120727.1_NLRP7; NP_789781.2_NLRP8; NP_127497.1_NLRP1; NP_001264057.1_LRRC31; NP_001263629.1_NLRP6; NP_006083.1_NOD1; NP_079101.4_PODNL1; NP_060110.4_CARMIL1

### Statistical analysis

Data expressed as mean ± SEM. Comparison between two groups was performed by a two-tailed *t*-test. A value of p<0.05 was considered to be statistically significant. All statistical analyses were calculated using Graph Pad Prism version 8.

### Materials availability

Plamisds and transgenic mice are available upon request. Please contact lead contact

## References

Allam, R., K.E. Lawlor, E.C. Yu, A.L. Mildenhall, D.M. Moujalled, R.S. Lewis, F. Ke, K.D. Mason, M.J. White, K.J. Stacey, A. Strasser, L.A. O’Reilly, W. Alexander, B.T. Kile, D.L. Vaux, and J.E. Vince. 2014. Mitochondrial apoptosis is dispensable for NLRP3 inflammasome activation but non apoptotic caspase 8 is required for inflammasome priming. Embo Rep. 15:982–990. doi:10.15252/embr.201438463.

Allam, R., M.H. Maillard, A. Tardivel, V. Chennupati, H. Bega, C.W. Yu, D. Velin, P. Schneider, and K.M. Maslowski. 2015. Epithelial NAIPs protect against colonic tumorigenesis. J Exp Med. 212:369–383. doi:10.1084/jem.20140474.

Arthur, J.S.C., and S.C. Ley. 2013. Mitogen-activated protein kinases in innate immunity. Nature Publishing Group. 1–14. doi:10.1038/nri3495.

Berg, D.F. van den, and A.A.T. Velde. 2020. Severe COVID-19: NLRP3 Inflammasome Dysregulated. Front Immunol. 11:1580. doi:10.3389/fimmu.2020.01580.

Broz, P., and V.M. Dixit. 2016. Inflammasomes: mechanism of assembly, regulation and signalling. Nat. Rev. Immunol. 1–14. doi:10.1038/nri.2016.58.

Broz, P., J. von Moltke, J.W. Jones, R.E. Vance, and D.M. Monack. 2010. Differential Requirement for Caspase-1 Autoproteolysis in Pathogen-Induced Cell Death and Cytokine Processing. Cell Host Microbe. 8:471–483. doi:10.1016/j.chom.2010.11.007.

Chennupati, V., D.F. Veiga, K.M. Maslowski, N. Andina, A. Tardivel, E.C.-W. Yu, M. Stilinovic, C. Simillion, M.A. Duchosal, M. Quadroni, I. Roberts, V.G. Sankaran, H.R. Macdonald, N. Fasel, A. Angelillo-Scherrer, P. Schneider, T. Hoang, and R. Allam. 2018. Ribonuclease inhibitor 1 regulates erythropoiesis by controlling GATA1 translation. J. Clin. Invest. 128:1597–1614. doi:10.1172/jci94956.

Desai, N., A. Neyaz, A. Szabolcs, A.R. Shih, J.H. Chen, V. Thapar, L.T. Nieman, A. Solovyov, A. Mehta, D.J. Lieb, A.S. Kulkarni, C. Jaicks, K.H. Xu, M.J. Raabe, C.J. Pinto, D. Juric, I. Chebib, R.B. Colvin, A.Y. Kim, R. Monroe, S.E. Warren, P. Danaher, J.W. Reeves, J. Gong, E.H. Rueckert, B.D. Greenbaum, N. Hacohen, S.M. Lagana, M.N. Rivera, L.M. Sholl, J.R. Stone, D.T. Ting, and V. Deshpande. 2020. Temporal and spatial heterogeneity of host response to SARS-CoV-2 pulmonary infection. Nature Communications. 1–15. doi:10.1038/s41467-020-20139-7.

Dickson, K.A., M.C. Haigis, and R.T. Raines. 2005. Ribonuclease inhibitor: structure and function. Prog. Nucleic Acid Res. Mol. Biol. 80:349–374. doi:10.1016/s0079-6603(05)80009-1.

Dickson, K.A., D.-K. Kang, Y.S. Kwon, J.C. Kim, P.A. Leland, B.-M. Kim, S.-I. Chang, and R.T. Raines. 2009. Ribonuclease inhibitor regulates neovascularization by human angiogenin. Biochemistry. 48:3804–3806. doi:10.1021/bi9005094.

Druilhe, A., S.M. Srinivasula, M. Razmara, M. Ahmad, and E.S. Alnemri. 2001. Regulation of IL-1beta generation by Pseudo-ICE and ICEBERG, two dominant negative caspase recruitment domain proteins. Cell Death Differ. 8:649–657. doi:10.1038/sj.cdd.4400881.

Ferreira, A.C., V.C. Soares, I.G. Azevedo-Quintanilha, S. da S.G. Dias, N. Fintelman-Rodrigues, C.Q. Sacramento, M. Mattos, C.S. Freitas, J.R. Temerozo, L. Teixeira, E.D. Hottz, E.A. Barreto, C.R.R. Pão, L. Palhinha, M. Miranda, D.C. Bou-Habib, F.A. Bozza, P.T. Bozza, and T.M.L. Souza. 2021. SARS-CoV-2 engages inflammasome and pyroptosis in human primary monocytes. Cell Death Discovery. 1–12. doi:10.1038/s41420-021-00428-w.

Furia, A., M. Moscato, G. Calì, E. Pizzo, E. Confalone, M.R. Amoroso, F. Esposito, L. Nitsch, and G. D’Alessio. 2011. The ribonuclease/angiogenin inhibitor is also present in mitochondria and nuclei. FEBS Lett. 585:613–617. doi:10.1016/j.febslet.2011.01.034.

Gong, T., L. Liu, W. Jiang, and R. Zhou. 2020. DAMP-sensing receptors in sterile inflammation and inflammatory diseases. Nat. Rev. Immunol. 1–18. doi:10.1038/s41577-019-0215-7.

Haigis, M.C., E.S. Haag, and R.T. Raines. 2002. Evolution of ribonuclease inhibitor by exon duplication. Mol. Biol. Evol. 19:959–963.

Henao-Mejia, J., E. Elinav, C.A. Thaiss, and R.A. Flavell. 2014. Inflammasomes and metabolic disease. Annu. Rev. Physiol. 76:57–78. doi:10.1146/annurev-physiol-021113-170324.

Hoffmann, M., H. Kleine-Weber, S. Schroeder, N. Krüger, T. Herrler, S. Erichsen, T.S. Schiergens, G. Herrler, N.-H. Wu, A. Nitsche, M.A. Müller, C. Drosten, and S. Pöhlmann. 2020. SARS-CoV-2 Cell Entry Depends on ACE2 and TMPRSS2 and Is Blocked by a Clinically Proven Protease Inhibitor. Cell. 181:271-280.e8. doi:10.1016/j.cell.2020.02.052.

Hoss, F., J.L. Mueller, F.R. Ringeling, J.F. Rodriguez-Alcazar, R. Brinkschulte, G. Seifert, R. Stahl, L. Broderick, C.D. Putnam, R.D. Kolodner, S. Canzar, M. Geyer, H.M. Hoffman, and E. Latz. 2019. Alternative splicing regulates stochastic NLRP3 activity. Nature Communications. 1–13. doi:10.1038/s41467-019-11076-1.

Huang, C., Y. Wang, X. Li, L. Ren, J. Zhao, Y. Hu, L. Zhang, G. Fan, J. Xu, X. Gu, Z. Cheng, T. Yu, J. Xia, Y. Wei, W. Wu, X. Xie, W. Yin, H. Li, M. Liu, Y. Xiao, H. Gao, L. Guo, J. Xie, G. Wang, R. Jiang, Z. Gao, Q. Jin, J. Wang, and B. Cao. 2020a. Clinical features of patients infected with 2019 novel coronavirus in Wuhan, China. Lancet. 395:497–506. doi:10.1016/s0140-6736(20)30183-5.

Huang, Q., X. Wu, X. Zheng, S. Luo, S. Xu, and J. Weng. 2020b. Targeting inflammation and cytokine storm in COVID-19. Pharmacol Res. 159:105051. doi:10.1016/j.phrs.2020.105051.

Jose, R.J., and A. Manuel. 2020. COVID-19 cytokine storm: the interplay between inflammation and coagulation. The Lancet Respiratory. 8:e46–e47. doi:10.1016/s2213-2600(20)30216-2.

Katoh, K., and D.M. Standley. 2013. MAFFT multiple sequence alignment software version 7: improvements in performance and usability. Mol. Biol. Evol. 30:772–780. doi:10.1093/molbev/mst010.

Kobe, B., and J. Deisenhofer. 1995. A structural basis of the interactions between leucine-rich repeats and protein ligands. Nature. 374:183–186. doi:10.1038/374183a0.

Kobe, B., and A.V. Kajava. 2001. 1-s2.0-S0959440X01002664-main. Curr. Opin. Struct. Biol. 11:1–8.

Kroemer, A., K. Khan, M. Plassmeyer, O. Alpan, M.A. Haseeb, R. Gupta, and T.M. Fishbein. 2020. Inflammasome activation and pyroptosis in lymphopenic liver patients with COVID-19. J Hepatol. 73:1258–1262. doi:10.1016/j.jhep.2020.06.034.

Kühn, R., F. Schwenk, M. Aguet, and K. Rajewsky. 1995. Inducible gene targeting in mice. Science. 269:1427–1429. doi:10.1126/science.7660125.

Lamkanfi, M., and V.M. Dixit. 2011. Inflammasomes and Their Roles in Health and Disease. Annu. Rev. Cell Dev. Biol. 28:120913150209003. doi:10.1146/annurev-cellbio-101011-155745.

Latz, E., T.S. Xiao, and A. Stutz. 2013. Activation and regulation of the inflammasomes. Nat. Rev. Immunol. 13:1–15. doi:10.1038/nri3452.

Lee, D.-J., F. Du, S.-W. Chen, M. Nakasaki, I. Rana, V.F.S. Shih, A. Hoffmann, and C. Jamora. 2015. Regulation and Function of the Caspase-1 in an Inflammatory Microenvironment. J. Invest. Dermatol. 135:2012–2020. doi:10.1038/jid.2015.119.

Letunic, I., and P. Bork. 2019. Interactive Tree Of Life (iTOL) v4: recent updates and new developments. Nucleic Acids Res. 47:W256–W259. doi:10.1093/nar/gkz239.

Li, J., H.L. Yin, and J. Yuan. 2008. Flightless-I regulates proinflammatory caspases by selectively modulating intracellular localization and caspase activity. J. Cell Biol. 181:321–333. doi:10.1083/jcb.200711082.

Li, P., H. Allen, S. Banerjee, S. Franklin, L. Herzog, C. Johnston, J. McDowell, M. Paskind, L. Rodman, and J. Salfeld. 1995. Mice deficient in IL-1 beta-converting enzyme are defective in production of mature IL-1 beta and resistant to endotoxic shock. Cell. 80:401–411. doi:10.1016/0092-8674(95)90490-5.

Liu, W., Y. Xie, J. Ma, X. Luo, P. Nie, Z. Zuo, U. Lahrmann, Q. Zhao, Y. Zheng, Y. Zhao, Y. Xue, and J. Ren. 2015. IBS: an illustrator for the presentation and visualization of biological sequences. Bioinformatics. 31:3359–3361. doi:10.1093/bioinformatics/btv362.

Lucas, C., P. Wong, J. Klein, T.B.R. Castro, J. Silva, M. Sundaram, M.K. Ellingson, T. Mao, J.E. Oh, B. Israelow, T. Takahashi, M. Tokuyama, P. Lu, A. Venkataraman, A. Park, S. Mohanty, H. Wang, A.L. Wyllie, C.B.F. Vogels, R. Earnest, S. Lapidus, I.M. Ott, A.J. Moore, M.C. Muenker, J.B. Fournier, M. Campbell, C.D. Odio, A. Casanovas-Massana, A. Obaid, A. Lu-Culligan, A. Nelson, A. Brito, A. Nunez, A. Martin, A. Watkins, B. Geng, C. Kalinich, C. Harden, C. Todeasa, C. Jensen, D. Kim, D. McDonald, D. Shepard, E. Courchaine, E.B. White, E. Song, E. Silva, E. Kudo, G. DeIuliis, H. Rahming, H.-J. Park, I. Matos, J. Nouws, J. Valdez, J. Fauver, J. Lim, K.-A. Rose, K. Anastasio, K. Brower, L. Glick, L. Sharma, L. Sewanan, L. Knaggs, M. Minasyan, M. Batsu, M. Petrone, M. Kuang, M. Nakahata, M. Campbell, M. Linehan, M.H. Askenase, M. Simonov, M. Smolgovsky, N. Sonnert, N. Naushad, P. Vijayakumar, R. Martinello, R. Datta, R. Handoko, S. Bermejo, S. Prophet, S. Bickerton, S. Velazquez, T. Alpert, T. Rice, W. Khoury-Hanold, X. Peng, Y. Yang, Y. Cao, Y. Strong, R. Herbst, A.C. Shaw, R. Medzhitov, W.L. Schulz, N.D. Grubaugh, C. Cruz, S. Farhadian, A.I. Ko, et al. 2020. Longitudinal analyses reveal immunological misfiring in severe COVID-19. Nature. 1–25. doi:10.1038/s41586-020-2588-y.

Lugrin, J., and F. Martinon. 2017. Detection of ASC Oligomerization by Western Blotting. Bio Protoc. 7:e2292–e2292. doi:10.21769/bioprotoc.2292.

Man, S.M., R. Karki, B. Briard, A. Burton, S. Gingras, S. Pelletier, and T.-D. Kanneganti. 2017. Differential roles of caspase-1 and caspase-11 in infection and inflammation. Nature Publishing Group. 1–11. doi:10.1038/srep45126.

Martinon, F., A. Mayor, and J. Tschopp. 2009. The inflammasomes: guardians of the body. Annu. Rev. Immunol. 27:229–265. doi:10.1146/annurev.immunol.021908.132715.

Martinon, F., V. Pétrilli, A. Mayor, A. Tardivel, and J. Tschopp. 2006. Gout-associated uric acid crystals activate the NALP3 inflammasome. Nature. 440:237–241. doi:10.1038/nature04516.

Matsushima, N., H. Miyashita, T. Mikami, and Y. Kuroki. 2010. A nested leucine rich repeat (LRR) domain: the precursor of LRRs is a ten or eleven residue motif. BMC Microbiol. 10:235–10. doi:10.1186/1471-2180-10-235.

Mehta, P., D.F. McAuley, M. Brown, E. Sanchez, R.S. Tattersall, J.J. Manson, and U.K.H.A.S. Collaboration. 2020. COVID-19: consider cytokine storm syndromes and immunosuppression. The Lancet. 395:1033–1034. doi:10.1016/s0140-6736(20)30628-0.

Miao, E.A., J.V. Rajan, and A. Aderem. 2011. Caspase-1-induced pyroptotic cell death. Immunol. Rev. 243:206–214. doi:10.1111/j.1600-065x.2011.01044.x.

Micco, A.D., G. Frera, J. Lugrin, Y. Jamilloux, E.-T. Hsu, A. Tardivel, A.D. Gassart, L. Zaffalon, B. Bujisic, S. Siegert, M. Quadroni, P. Broz, T. Henry, C.A. Hrycyna, and F. Martinon. 2016. AIM2 inflammasome is activated by pharmacological disruption of nuclear envelope integrity. Proceedings of the National Academy of Sciences. 113:E4671–80. doi:10.1073/pnas.1602419113.

Monti, D.M., N.M. Gesualdi, J. Matousek, F. Esposito, and G. D’Alessio. 2007. The cytosolic ribonuclease inhibitor contributes to intracellular redox homeostasis. FEBS Lett. 581:930–934. doi:10.1016/j.febslet.2007.01.072.

Ng, A.C.Y., J.M. Eisenberg, R.J.W. Heath, A. Huett, C.M. Robinson, G.J. Nau, and R.J. Xavier. 2011. Human leucine-rich repeat proteins: a genome-wide bioinformatic categorization and functional analysis in innate immunity. Proc. Natl. Acad. Sci. U.S.A. 108 Suppl 1:4631–4638. doi:10.1073/pnas.1000093107.

Nie, J., Q. Li, J. Wu, C. Zhao, H. Hao, H. Liu, L. Zhang, L. Nie, H. Qin, M. Wang, Q. Lu, X. Li, Q. Sun, J. Liu, C. Fan, W. Huang, M. Xu, and Y. Wang. 2020. Establishment and validation of a pseudovirus neutralization assay for SARS-CoV-2. Emerging Microbes & Infections. 9:680–686. doi:10.1080/22221751.2020.1743767.

Papin, S., S. Cuenin, L. Agostini, F. Martinon, S. Werner, H.-D. Beer, C. Grütter, M. Grütter, and J. Tschopp. 2007. The SPRY domain of Pyrin, mutated in familial Mediterranean fever patients, interacts with inflammasome components and inhibits proIL-1beta processing. Cell Death Differ. 14:1457–1466. doi:10.1038/sj.cdd.4402142.

Pedraza-Alva, G., L. Pérez-Martínez, L. Valdez-Hernández, K.F. Meza-Sosa, and M. Ando-Kuri. 2015. Negative regulation of the inflammasome: keeping inflammation under control. Immunol. Rev. 265:231–257. doi:10.1111/imr.12294.

Rodrigues, T.S., K.S.G. de Sá, A.Y. Ishimoto, A. Becerra, S. Oliveira, L. Almeida, A.V. Goncalves, D.B. Perucello, W.A. Andrade, R. Castro, F.P. Veras, J.E. Toller-Kawahisa, D.C. Nascimento, M.H.F. de Lima, C.M.S. Silva, D.B. Caetite, R.B. Martins, I.A. Castro, M.C. Pontelli, F.C. de Barros, N.B. do Amaral, M.C. Giannini, L.P. Bonjorno, M.I.F. Lopes, R.C. Santana, F.C. Vilar, M. Auxiliadora-Martins, R. Luppino-Assad, S.C.L. de Almeida, F.R. de Oliveira, S.S. Batah, L. Siyuan, M.N. Benatti, T.M. Cunha, J.C. Alves-Filho, F.Q. Cunha, L.D. Cunha, F.G. Frantz, T. Kohlsdorf, A.T. Fabro, E. Arruda, R.D.R. de Oliveira, P. Louzada-Junior, and D.S. Zamboni. 2021. Inflammasomes are activated in response to SARS-CoV-2 infection and are associated with COVID-19 severity in patients. Journal of Experimental Medicine. 218. doi:10.1084/jem.20201707.

Schroder, K., and J. Tschopp. 2010. The Inflammasomes. Cell. 140:821–832. doi:10.1016/j.cell.2010.01.040.

Spinetti, T., C. Hirzel, M. Fux, L.N. Walti, P. Schober, F. Stueber, M.M. Luedi, and J.C. Schefold. 2020. Reduced Monocytic Human Leukocyte Antigen-DR Expression Indicates Immunosuppression in Critically Ill COVID-19 Patients. Anesthesia & Analgesia. 131:993–999. doi:10.1213/ane.0000000000005044.

Strowig, T., J. Henao-Mejia, E. Elinav, and R. Flavell. 2012. Inflammasomes in health and disease. Nature. 481:278–286. doi:10.1038/nature10759.

Stutz, A., G.L. Horvath, B.G. Monks, and E. Latz. 2013. ASC speck formation as a readout for inflammasome activation. Methods Mol. Biol. 1040:91–101. doi:10.1007/978-1-62703-523-1_8.

Tavernier, Q., E. Bennana, V. Poindessous, C. Schaeffer, L. Rampoldi, N. Pietrancosta, and N. Pallet. 2018. Regulation of IRE1 RNase activity by the Ribonuclease inhibitor 1 (RNH1). Cell Cycle. 17:1901–1916. doi:10.1080/15384101.2018.1506655.

Toldo, S., R. Bussani, V. Nuzzi, A. Bonaventura, A.G. Mauro, A. Cannatà, R. Pillappa, G. Sinagra, P. Nana-Sinkam, P. Sime, and A. Abbate. 2020. Inflammasome formation in the lungs of patients with fatal COVID-19. Inflamm. Res. 1–4. doi:10.1007/s00011-020-01413-2.

Walle, L.V., M. Lamkanfi, and P. Vandenabeele. 2008. The mitochondrial serine protease HtrA2/Omi: an overview. Cell Death Differ. 15:453–460. doi:10.1038/sj.cdd.4402291.

Yamasaki, S., P. Ivanov, G.-F. Hu, and P. Anderson. 2009. Angiogenin cleaves tRNA and promotes stress-induced translational repression. J. Cell Biol. 185:35–42. doi:10.1083/jcb.200811106.

Zhou, R., A.S. Yazdi, P. Menu, and J. Tschopp. 2010. A role for mitochondria in NLRP3 inflammasome activation. Nature. 1–6. doi:10.1038/nature09663.

