## Supplementary material for "LRR protein RNH1 dampens the inflammasome activation and is associated with adverse clinical outcomes in COVID-19 patients": Supplemetary figures

**Figure S1- S5**

### Figure S1

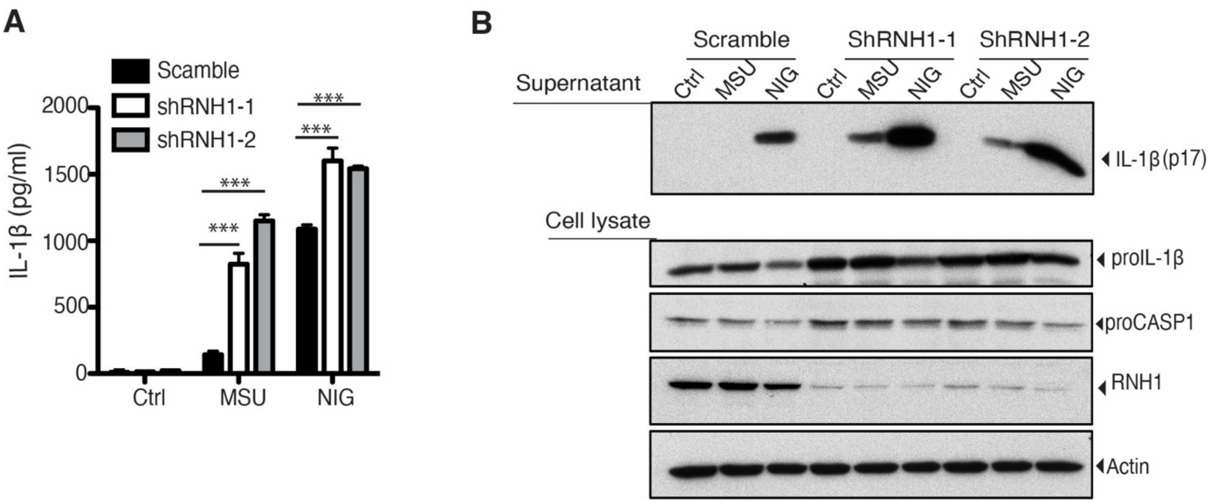


**Fig S1. RNH1 knock down aggravated NLRP3 inflammasome activation. (A** and **B)** PMA differentiated scramble and RNH1 knock down THP1 cells were stimulated with Nigericin (NIG) (5 µM) for 1h and for 5 h with MSU (500 µg). (**A**) Supernatant were collected and IL-1β ELISA was performed. Data are means ± SEM of pooled data from three independent experiments. Statistical analyses were performed using a two-tailed *t*-test. (**B**) Cell lysates and supernatants were analysed for pro- and cleaved- forms of IL-1β by Western blot. Blots were representative of two independent experiments.

### Figure S2

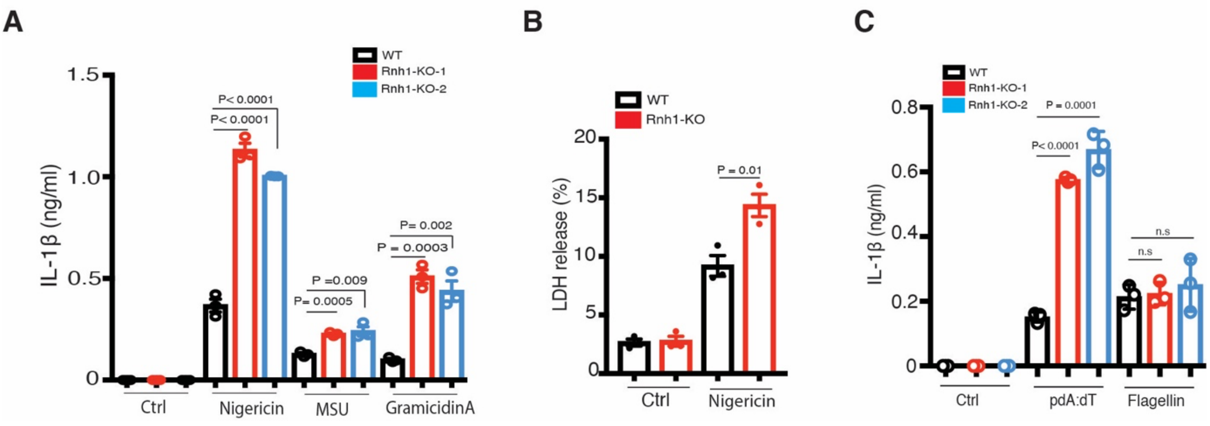


**Fig S2. Loss of RNH1 increased inflammasome activation in iMAC cells**

**(A)** WT and Rnh1-KO iMAC cells were primed with 500 ng of LPS for 3h and then stimulated with Nigericin (5µM) for 1h or with MSU (500µg) and Gramicidin A (30 µg) for 5h. Supernatant were collected and IL-1β ELISA was performed. (**B**) WT and Rnh1-KO iMAC cells were primed with 500 ng of LPS for 3h and then stimulated with Nigericin (5µM) for 8h. Supernatants were collected, and cell death was measured by LDH assay. (**C**) iMAC cells were primed with 500 ng of LPS for 3h and then transfected with AIM2 agonist poly dA:dT (5 µg) or NAIP/NLRC4 agonist cytosolic flagellin (600 ng) for 5h. Supernatants were collected and IL-1β ELISA was performed . All the data are means ± SEM of pooled data from three independent experiments. Statistical analyses were performed using a two-tailed *t*-test.

### Figure S3

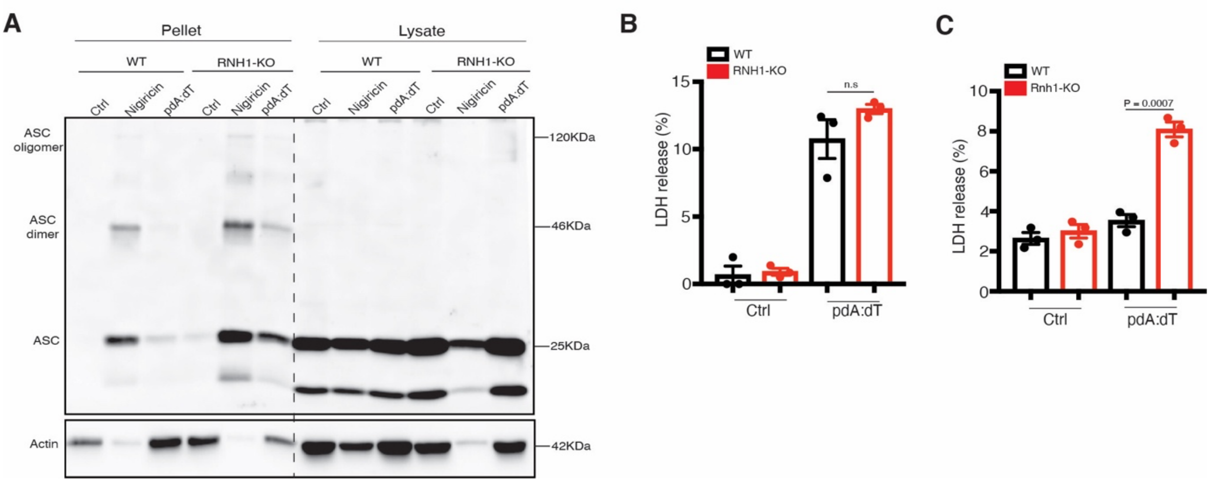


**Fig S3. Loss of RNH1 increased ASC oligomerization. (A**) PMA differentiated WT and RNH1-KO THP1 were treated with nigericin (5µM) for 1h and poly(dA:dT) (5 µg) for 5h. Cross-linked pellets (Pellets) or soluble lysates (Lysates) were immunoblotted for ASC. Blots are representative of three independent experiments. (**B** and **C)** WT and RNH1-KO THP1 and iMAC cells were stimulated with AIM2 agonist poly dA:dT (5 µg) for 18h. Supernatant were collected and cell death was measured by LDH assay Data are means ± SEM of pooled data from three independent experiments. Statistical analyses were performed using a two-tailed *t*-test.

**Figure S4**

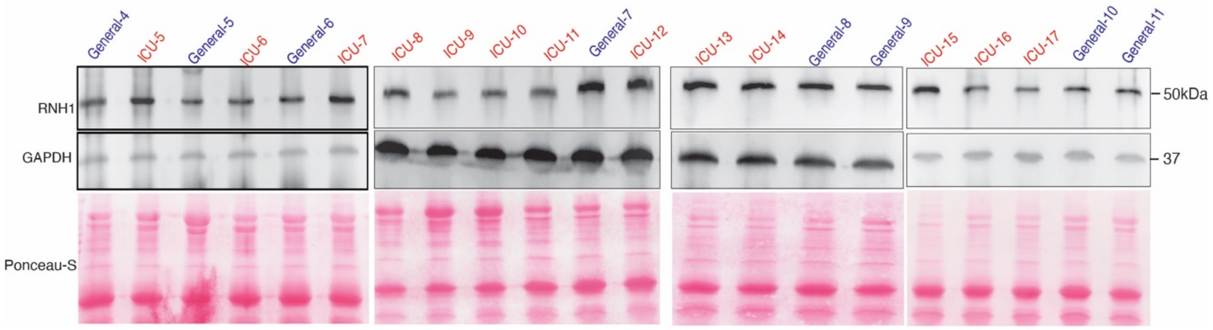


**Fig S4. RNH1 expression negatively correlates with diseases severity in COVID-19 patients.** Total cell lysates from peripheral blood buffy coats of ICU and general ward admitted COVID-19 patients were analysed for RNH1 protein levels by Western blot. Blots were repeated for three times.

**Figure S5**

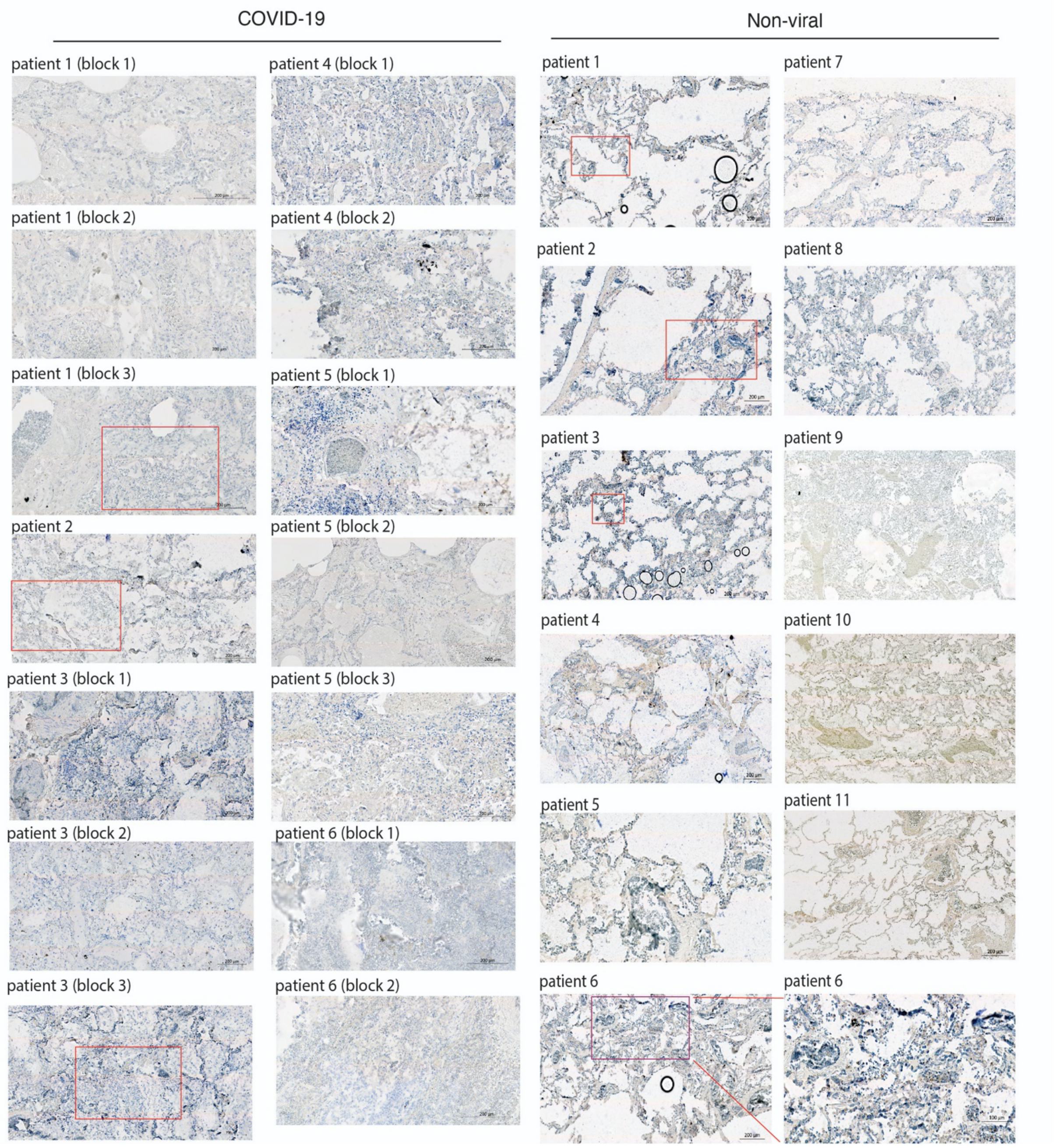


**Fig S5. RNH1 expression is decreased in COVID-19 patients lung biopsies compared to non-viral patients.** Postmortem lung tissue from deceased persons with either COVID-19 (n =8) or non-viral (n= 13) causes of death were stained for RNH1 and imaged using a Zeiss axioscan Z1. Subsequent image analysis was performed using Zeiss ZEN software to extract images at different magnifications. Insets show higher magnification of area indicated in the red boxes. Brown staining indicates RNH1 positive cells.
