## Supplementary material for "LRR protein RNH1 dampens the inflammasome activation and is associated with adverse clinical outcomes in COVID-19 patients": Supplemetary Table-1

| **Table S1:** Baseline demographics, disease severity, and clinical outcome. | | | | | |
| --- | --- | --- | --- | --- | --- |
|  | | **ICU patients with COVID-19**  (n=17) | **Hospitalized COVID-19 patients**  (n=11) | **Total cohort** (n=28) | between group  p-value |
| Demographics | - Age (years) | 69.6 [66.9 , 77.2] | 55 [49 , 70.5] | 68.8 [54.95 , 73.33] | 0.082 |
|  | - Gender (male, %) | 14 (82.4) | 9 (81.8) | 23 (82.1) | 1 |
|  | - Body Mass Index | 26.2 [25.25 , 29.98] | 27.3 [24.44 , 29.22] | 26.6 [24.89 , 29.79] | 0.957 |
|  | - APACHE-II score (first 24 hours) | 21 [19 , 26] | - | - | - |
|  | - SAPS II score (first 24 hours) | 45 [42 , 59] | - | - | - |
|  | - SOFA score (baseline) | 8 [7 , 10] | - | - | - |
| Comorbidity data | - Charleson Comorbidity Index   (total score) | 4 [3 , 7] | - | - | - |
|  | - Myocardial infarction (No./%) | 4 (23.53) | - | - | - |
|  | - Chronic heart failure (No./%) | 1 (5.88) | - | - | - |
|  | - Peripheral vascular disease (No./%) | 1 (5.88) | - | - | - |
|  | - Cerebrovascular accident (No./%) | 2 (11.76) | - | - | - |
|  | - Dementia (No./%) | 0 (0) | - | - | - |
|  | - COPD (No./%) | 2 (11.76) | - | - | - |
|  | - Connective tissue disease (No./%) | 0 (0) | - | - | - |
|  | - Peptic ulcer disease (No./%) | 0 (0) | - | - | - |
|  | - Liver disease (0-3) (No./%) | 2 (11.76) | - | - | - |
|  | - Diabetes (0-2) (No./%) | 7 (41.18) | - | - | - |
|  | - Hemiplegia (No./%) | 0 (0) | - | - | - |
|  | - Moderate to severe CKD (No./%) | 2 (11.76) | - | - | - |
|  | - Solid tumor (0-6) (No./%) | 1 (5.88) | - | - | - |
|  | - Leukemia (No./%) | 0 (0) | - | - | - |
|  | - Lymphoma (No./%) | 1 (5.88) | - | - | - |
|  | - HIV/ AIDS (No./%) | 1 (5.88) | - | - | - |
| Laboratory data | - C-reactive protein (mg/L) | 148 [87 , 315] | 43 [12.5 , 101.5] | 100.5 [54.25 , 196.5] | 0.001 |
|  | - Procalcitonin levels (ng/ml) | 1.1 [0.35 , 9.82] | 0.2 [0.15 , 0.24] | 0.4 [0.25 , 5] | 0.004 |
|  | - Total leukocyte count (G/L) | 8.4 [6.98 , 10.5] | 6.1 [4.82 , 6.88] | 7 [5.95 , 9.94] | 0.004 |
|  | - Total lymphocyte count (G/L) | 0.6 [0.44 , 0.77] | 1.1 [0.88 , 1.36] | 0.7 [0.51 , 1.12] | 0.1 |
|  | - Platelet count (G/L) | 208 [117 , 225] | 203 [153 , 247] | 205.5 [147.25 , 231.25] | 0.572 |
|  | - Serum potassium (mmol/L) | 4.2 [4 , 4.6] | 3.8 [3.7 , 3.9] | 4 [3.9 , 4.2] | 0 |
|  | - Serum creatinine (µmol/L) | 103 [86 , 160] | 73 [57.5 , 86.5] | 89 [68 , 132.25] | 0.008 |
|  | - D-Dimers (µg/L) | 1291 [1183 , 1447] | 490 [459 , 1276] | 1262 [1017 , 1479.5] | 0.482 |
| Follow-up | - Days on ICU | 14 [8 , 21] | - | - | - |
|  | - Days in hospital | 17 [9 , 24] | 6 [3.5 , 8] | 10.5 [5.75 , 21.75] | 0.004 |
|  | - Days on antibiotics | 10 [8 , 14] | 0.5 [0 , 4] | 8 [3 , 14] | 0.001 |
|  | - Total days on mechanical ventilation | 15 [9 , 20.56] | - | - | - |
|  | - Renal replacement at any time   (No./%) | 9 (52.9) | - | - | - |
|  | - On vasopressors at any time   (No./%) | 17 (100) | - | - | - |
|  | - Norepinephrine dose   (cumulative dose/ ICU days; mg) | 2.9 [1.9 , 5.77] | - | - | - |
|  | - ICU Mortality (No./%) | 6 (35.3) | - | - | - |
|  | - Hospital mortality (No./%) | 6 (35.3) | 0 (0) | 6 (21.4) |  |
| Demographical data, baseline comorbidities, laboratory data, and clinical follow-up is given for patients with primary admission to ICU vs. normal ward. G= Giga, L= Liters, APACHE-II= Acute Physiology and Chronic Health Evaluation- II score, SAPS-2= Simplified Acute Physiology Score-2, SOFA= Sepsis-related organ failure assessment score, COPD= chronic obstructive pulmonary disease, HIV= human immunodeficiency virus, AIDS= acquired immunodeficiency syndrome. Numbers (No.) with percentages are given, as indicated. Continuous data are reported as median [quartiles]. Between group p-values from Mann-Whitney U tests and Fisher’s exact tests are given for ICU vs. non-ICU (normal ward) populations. Between group p-values are given for ICU vs. non-ICU (normal ward) populations. | | | | | |

**Table S1: Patient demographics, disease severity, and clinical outcomes**

Demographical data, baseline comorbidities, laboratory data, and clinical follow-up is given for patients with primary admission to ICU vs. normal ward. G= Giga, L= Liters, APACHE-II= Acute Physiology and Chronic Health Evaluation- II score, SAPS-2= Simplified Acute Physiology Score-2, SOFA= Sepsis-related organ failure assessment score, COPD= chronic obstructive pulmonary disease, HIV= human immunodeficiency virus, AIDS= acquired immunodeficiency syndrome. Numbers (No.) with percentages are given, as indicated. Continuous data are reported as median [quartiles]. Between group p-values from Mann-Whitney U tests and Fisher’s exact tests are given for ICU vs. non-ICU (normal ward) populations. Between group p-values are given for ICU vs. non-ICU (normal ward) populations.
